# Effect of human urinary microenvironment and fluid flow on antibiotic and phage therapy efficacy against uropathogenic *Escherichia coli*

**DOI:** 10.1101/2025.11.12.688052

**Authors:** Ramon Garcia Maset, Aaron Crowther, Victoria Chu, Davide Di Grande, Rizka O.A. Jariah, Jenny Yoon, Dalia Blumgart, Marloes A.E Hodek, Nicholas Yuen, Benjamin O. Murray, Laia Pasquina-Lemonche, Sara N.H Alghamd, Andrew D. Millard, Gareth LuTheryn, Melissa E.K. Haines, Martha R.J. Clokie, Eleanor Stride, Dario Carugo, Jennifer L. Rohn

**Author notes:** Co-senior authors (Dario Carugo, Jennifer L. Rohn).

## Abstract

Urinary tract infections (UTI) remain a major global health burden, with high recurrence despite antibiotic treatment. The escalating prevalence of antimicrobial resistance further compromises therapeutic efficacy, contributing to an estimated 260,000 deaths annually. Conventional *in vitro* susceptibility assays often fail to predict clinical outcomes, underscoring the urgent need for physiologically relevant infection models.

Here, we examined how microenvironmental complexity shapes uropathogenic *Escherichia coli* (UPEC) responses to antibiotics and bacteriophages using: human urine, a three-dimensional urothelial microtissue model (3D-UHU), and a novel mesofluidic system (P-FLO) that introduces physiologically relevant flow dynamics to the 3D-UHU. P-FLO was engineered from cost-effective 3D-printed components compatible with standard Transwell systems.

Among the antibiotics tested, nitrofurantoin exhibited the greatest potency in minimum inhibitory concentration assays, but it failed to fully eradicate infection within the more physiological 3D-UHU model. A bacteriophage cocktail (LCPR1) showed markedly reduced activity in urine compared with nutrient-rich media, highlighting the influence of infection-site conditions. In contrast, in 3D-UHU, LCRP1 modulated host responses without reducing bacterial burden. Combination therapy (nitrofurantoin + LCPR1) eliminated planktonic bacteria under static conditions but offered no added benefit against adherent or intracellular populations relative to antibiotic monotherapy. Incorporating flow revealed additional layers of complexity, where shear stress induced bacterial elongation and attachment and altered drug performance, diminishing the efficacy of nitrofurantoin and combination therapy against planktonic populations despite increased drug exposure.

Together, these findings demonstrate that the bladder microenvironment and its mechanical forces modulate host-pathogen interactions and profoundly influence UPEC infection dynamics and therapeutic outcomes, emphasizing the need for advanced, physiologically informed models to guide treatment strategies in the post-antibiotic era.

## Introduction

With 400 million cases per year, urinary tract infections (UTI) are among the most prevalent human infections, disproportionately affecting women and contributing significantly to the global burden of the antimicrobial resistance (AMR) crisis (with ∼260,000 AMR-associated UTI deaths annually)^1,2^. Approximately 80% of UTIs are caused by uropathogenic *E. coli* (UPEC), which is also linked to the highest number of deaths associated with AMR^3–5^. Although UTI is often regarded as uncomplicated and readily treatable, its high recurrence rate belies this view, with the biological and ecological processes governing infection, clearance and treatment failure poorly understood^6^.

The urothelium is a highly dynamic environment, shaped by fluctuating osmolarity, pH, nutrient composition, urine flow, and stretching cycles during the filling and voiding phases^7^. These factors influence bacterial physiology (adhesion, filamentation, invasion of uroepithelial cells, and biofilm formation), while host-derived components such as urea concentration, antimicrobial peptides, immune cell infiltration and the resident urinary microbiota further modulate pathogen physiology and antibiotic susceptibility^8–11^. Together, the urinary tract microenvironment creates a complex niche that modulates bacterial behaviour and influences the host/pathogen interaction alongside treatment efficacy.

Despite the availability of effective antibiotics, recurrence and relapse are common (up to 30% of cases), and repeated antimicrobial exposure contributes to the escalating global problem of AMR^12,13^. Treatment failure in UTI often occurs even when pathogens appear to be susceptible to conventional drugs. Standard *in vitro* susceptibility assays are typically conducted under nutrient-rich and static conditions that fail to recapitulate the physicochemical and fluid mechanical complexity of the urinary tract^14^. As a result, antimicrobial efficacy measured by minimum inhibitory concentration (MIC) often correlates poorly with clinical outcomes^15,16^.

Existing experimental models of UTI (ranging from cell culture to murine infection systems) have provided important mechanistic insights into UPEC behaviour but exhibit significant translational limitations. Murine models, for example, differ from humans in terms of bladder physiology, immune response, and urine composition^17^. Furthermore, pharmacokinetic and pharmacodynamic parameters of antibiotics often translate poorly between mice and humans, hindering efforts to define optimal treatment regimens^18^. Clinical recommendations for antibiotic choice, dosing, and duration thus remain largely empirical and lack robust evidence for efficacy in diverse patient populations^19^.

Recent tissue-engineering advances have provided human-relevant models to study UTI pathology. For example, a donor-derived human bladder organoid model has been developed to investigate UPEC and bacteriophage interaction at the molecular level^20^. To modulate stretching patterns during micturition and immune cell infiltration, a bladder-on-chip model was established to investigate intracellular reservoirs^21^. We recently developed the static <u>3</u>-dimensional <u>u</u>rine-tolerant <u>h</u>uman <u>u</u>rothelial microtissue Transwell model (3D-UHU), which allows investigation of UTI in a fully stratified and differentiated urothelium in a 100% human urine apical environment^22,23^. Together, these innovations and others are driving a new era of personalized, mechanism-based approaches to investigate UTI and antimicrobial therapies.

Treatment failure in UTI underscores the urgent need to explore alternative or adjunctive therapeutic approaches that can overcome bacterial adaptation. Increasing evidence suggests that non-genetic drivers including persistence, intracellular reservoirs and biofilm formation enable UPEC and other uropathogens to withstand otherwise effective antibiotic therapy. Innovative strategies that exploit different antibacterial mechanisms or synergise with existing antibiotics are gaining renewed attention. Bacteriophage (phage) therapy, either alone or in combination with antibiotics, offers a promising avenue to address these challenges^24^. Phages exhibit high specificity, self-amplifying dynamics and, in some cases, the capacity to disrupt biofilms and target metabolically quiescent bacterial populations^24^. Recent studies demonstrate that phage-antibiotic combinations can enhance bacterial clearance, suppress resistance evolution, and restore antibiotic susceptibility *in vitro* and in animal models^25^. Yet how such interventions perform within the complex urinary microenvironment and how host factors, such as flow dynamics and urine composition, influence phage activity remain largely unexplored.

Here, we investigated the antimicrobial efficacy of the common antibiotic nitrofurantoin against UPEC in increasingly complex environments. These included human urine; the 3D-UHU human urothelial static microtissue model; and the <u>P</u>lug-<u>F</u>low <u>L</u>inked <u>O</u>rganoid (P-FLO) system, a novel mesofluidic version of 3D-UHU developed from inexpensive, simple 3D-printed components allowing us to adopt commercially available Transwells to create a more physiologically relevant flow dynamic environment. We also used the static model as well as P-FLO to assess the activity of bacteriophages alone and in combination with antibiotics. These experiments demonstrated the marked effect of microenvironment on the response of UPEC to infection and treatment.

## Materials and methods

### Bacterial strains and bacterial growth for bladder microtissue infections

Uropathogenic *Escherichia coli* (UPEC) strains used in this study included CFT073 (pyelonephritis isolate) and HM50 (asymptomatic bacteriuria isolate), obtained from the American Type Culture Collection (ATCC). UTI89 isolated from patients with acute cystitis and its GFP-expressing derivative (UTI89-GFP), were generously provided by Scott Hultgren’s lab (Washington University in St. Louis, USA). UTI89-RPF^26^ was provided by Molly Ingersoll’s lab (Institut Cochin, Paris). The non-pathogenic laboratory strain *E. coli* MG1655 was kindly provided by Meriem El Karoui (University of Edinburgh, UK). The clinical isolates from Royal Free Hospital (London, UK) include three previously described UPEC strains isolated from clean-catch midstream urine samples of patients with chronic UTI (ClinA, EC3 and EC4), and two commensal *E. coli* isolates (COM1 to COM2) similarly recovered from healthy individuals^27^. Bacterial strains were routinely plated on LB agar (BD Difco LB agar, Lennox) and supplemented with appropriate antibiotics (kanamycin 50 μg ml^−1^ or ampicillin 100 μg ml^−1^) for the GFP and RFP expression, respectively.

### LCPR1-Phage cocktail propagation and purification

Host liquid propagation was used for each phage with its respective host (**Table 1**). In brief, bacterial strains were cultured to log phase (OD_600_ ∼ 0.15-0.2) using an overnight culture as a starting point. 200 μl of bacterial culture and 200 μl of phage stock were then added to 20 mL of LB media and incubated in a shaking incubator at 37 °C overnight (∼18 h). The next day, the lysate was centrifuged at 4,000 × g for 10 min and the supernatant was filter-sterilised through 0.2 μm pore-sized filters (Fisherbrand™). The phage stock titre was determined using plaque assays as described previously^28^ and phage stocks were stored at 4°C. The phage cocktail was produced by mixing all 9 phages in a 1:1 ratio (10^11^ PFU mL^-1^ each of phage stock), then diluted for the final LCPR1 cocktail to a concentration of 10^10^ PFU mL^-1^. The cocktail was then run through an Endotrap HD column (LIONEX, GmBH) following the manufacturer’s instructions. In brief, 1 mL of purified phage in SM-Buffer was run through the column using gravity. 200 μl of void volume was collected in a separate tube.

**Table 1.**
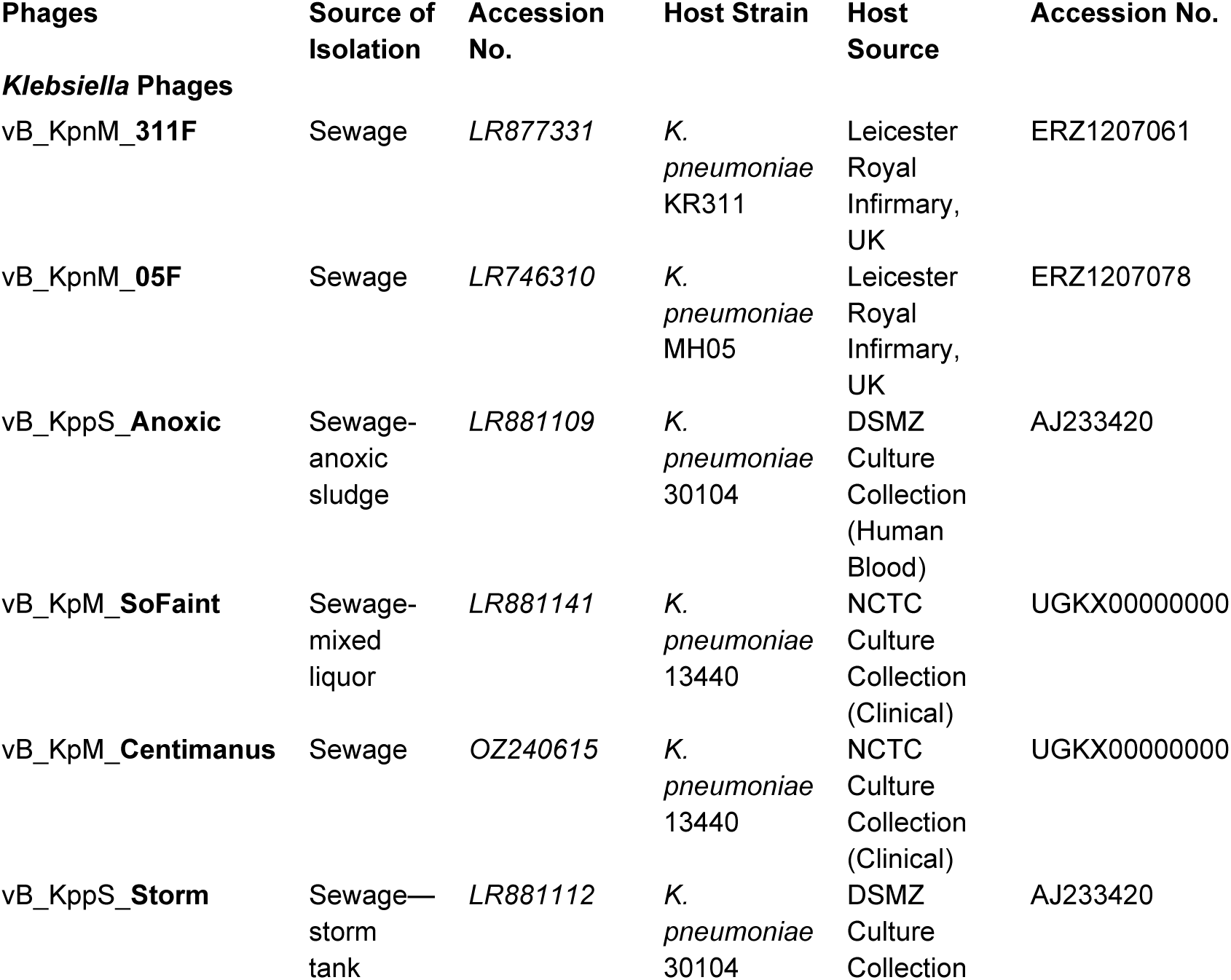
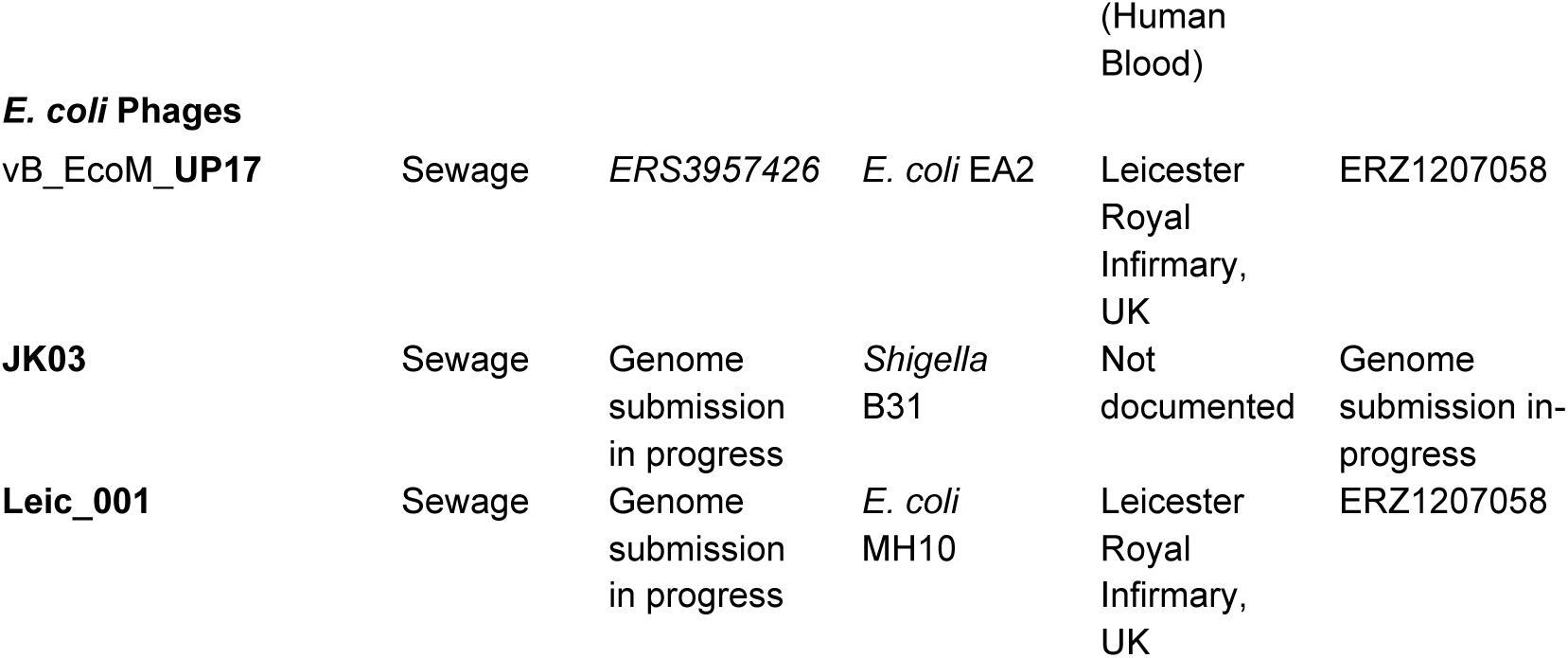
Phages included in LCPR1 cocktail and original propagation hosts used in this study.

### Bacterial growth curves

Bacterial cultures were made by inoculating a single colony from fresh agar plates into 5 mL of LB broth (Sigma-Aldrich) and placed in an orbital shaking incubator at 37°C for 24 h at 200 rpm. Cultures were adjusted to OD_600_ ∼ 0.1 in phosphate-buffered saline (PBS, Gibco) and further diluted 1:100 in cation-adjusted Müller-Hinton broth (caMHB, Sigma Aldrich); 100% pooled human urine (pHU); or a mixture of the two (70%pHU:30%caMHB). The pHU was obtained commercially (BioIVT) or obtained and decellularized from healthy volunteers of both sexes under ethics granted by the UCL Research Ethics Committee (Project ID 23285/002). Each batch of pHU, whether commercially sourced or collected, consisted of samples from 10/15 mixed sex donors. Then, 200 μL were transferred to flat-bottomed 96-well plates (Thermofisher) in triplicate technical replicates, and growth curves were generated using a microtiter plate reader (Tecan Spark® 10 M), measuring OD_600_ every 30 mins over 16 h at 37°C. Three biological experiments were performed with three technical replicates per experiment.

### Minimum inhibitory concentration (MIC)

MIC was determined according to the standard Clinical Laboratory Standards Institute (CLSI) broth microdilution method (M07-A9-2012)^29^ using caMHB. Additionally, MICs were determined using pHU mixed in caMHB (70:30 v/v%). Overnight cultures of bacteria were grown in caMHB and in 70% pHU in shaking conditions at 37 °C. Bacterial cultures were adjusted to OD_600_ ∼ 0.08-0.1 in PBS, corresponding to a bacterial concentration of ∼10^8^ colony forming units per mL (CFU mL^-1^). The solution was diluted 100-fold to obtain a concentration of 10^6^ CFU mL^-1^ in caMHB or 70% pHU. Antibiotics were dissolved in camHB or 70% pHU, and 100 μL of each antibiotic solution was added and serially diluted into the 96-well plates followed by the addition of the same volume of bacterial suspension, resulting in a final bacterial density of 5×10^5^ CFU mL^-1^. The microwell plates were incubated at 37 °C statically for 24 h, and growth was evaluated by eye. Triplicates were performed for each concentration. Bacteria alone or uninfected media were used as positive and negative controls, respectively. The experiment was repeated three times, and the highest MIC value obtained was reported.

### Minimum bactericidal concentration (MBC) and dose-response

After MIC determination, the bactericidal effect of the antibiotics was evaluated by adding 20 μL of resazurin sodium dye (Cambridge Bioscience, UK) with a final concentration of 0.5 mg mL^−1^ per well^30^. The plates were then incubated for 6 h at 37 °C. A noticeable change of colour could be observed for bacterial cells exhibiting growth (pink) versus non-detectable growth (blue). The lowest concentration where blue colour was observed (bactericidal concentration) was recorded as the MBC. Three independent experiments were performed. After the incubation of resazurin, the fluorescence signal in the 96-well plate was measured in a Tecan Spark® 10 M plate reader (excitation wavelength = 571 nm; emission wavelength = 584 nm). The bacterial viability in each well was then calculated using the positive control (untreated bacteria) and the negative control or blank (sterile media with resazurin) as follows:

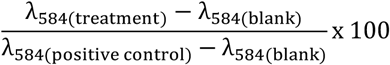

### Time-killing assay

Overnight cultures in caMHB and 70% pHU:caMHB were grown at 37°C under shaking conditions (200 rpm). These cultures were subsequently diluted to OD_600_ ∼ 0.01 in their respective media and grown to mid-exponential growth phase (OD_600_ ∼ 0.3), for approximately 2-3h. Bacterial cultures were adjusted to OD_600_ ∼ 0.08-0.1 in PBS. The bacterial solution was diluted 100-fold to obtain a concentration of 10^6^ CFU mL^-1^ in a total volume of 5 mL of caMHB or 70% pHU. Nitrofurantoin was added at the desired concentrations, and cultures were then placed in the shaking incubator at 37°C. Samples (80 μL) were taken at several time points as shown on the resulting graphs. Serial dilutions were performed with phosphate-buffered saline (PBS) in 96-well plates and plated on LB agar plates for CFU enumeration. Three independent experiments were performed. Untreated bacteria were used as a positive control.

### Human bladder cells and the 3D-UHU urothelial microtissue 3D model

Spontaneously immortalized, non-transformed human bladder epithelial cells (HBLAK; CELLnTEC) were cultured as described previously^22^. Briefly, HBLAK were cultured in CnT-Prime medium (CELLnTEC) under standard conditions (37°C, 5% CO₂, humidified atmosphere). Cells at 80–90% confluence were passaged by first rinsing with calcium- and magnesium-free phosphate-buffered saline (PBS; Gibco), followed by enzymatic detachment using CnT-Accutase solution (CELLnTEC) for 10 minutes at 37°C. Three-dimensional urine-tolerant human urothelial models (3D-UHU) were created as described previously^22^. Briefly, between passages 9 and 13, HBLAK cells at 80–90% confluency were enzymatically detached and resuspended at a density of 5 × 10⁵ cells mL^-1^ in pre-warmed CnT-Prime Epithelial Proliferation medium (CnT-PR, CELLnTEC). 400 μL of cell suspension were seeded onto 12-mm diameter, 0.4-μm pore polycarbonate membranes in plastic Transwell inserts (Corning), with 1.5 mL of medium added to the basal compartment. After 48 h of incubation at 37°C and 5% CO₂, the medium in both apical and basal compartments was replaced with CnT-Prime Epithelial 3D Airlift Medium (CnT-PR-3D, CELLnTEC), designated as day 0. Following 48h infection incubation, the Tranwells were transferred onto 12-well Thincert plates (Greiner) and the apical medium was substituted with 300 μL of sterile-filtered pooled human urine (BioIVT), collected from 10/15 healthy donors of both sexes, while fresh differentiation medium was replenished in the basal chamber (4.5 mL). Urine and medium were refreshed every 3-4 days until days 16-20, at which point the matured urothelial microtissue models were used for infections.

### Antimicrobial efficacy in the 3D-UHU model under static conditions

Fully differentiated 3D-UHU microtissue models from day 16 were used for the infection. Overnight cultures of *E. coli* UTI89 (LB) and *E. coli* UTI89-GFP (LB + supplemented with 50 μg mL^-1^ of kanamycin) were cultured in static conditions over 2 days (∼48h). Bacterial concentration (bacterial cells mL^-1^) was quantified with the QUANTOM Tx Microbial Cell Counter (Logos) following the manufacturer’s protocol. Then, the bacterial overnight culture was resuspended in human urine at 5.0 x 10^6^ bacterial cells mL^-1^ (similar inoculum as the MIC assay). Then, the models were infected for 12h at 37°C with 5% CO_2_. After 12h, two doses of antibiotic (nitrofurantoin, 256 μg mL^-1^), phage cocktail (LCPR1, 1 x 10^9^ phage mL^-1^), and the combination of both treatments (256 μg mL^-1^ and 1 x 10^9^ phage mL^-1^) were added in intervals of 4 h. Before and after the treatment addition, the urine in the apical compartment was collected and serially diluted for CFU enumeration in LB agar plates. At the end of the second treatment, urine was collected for CFU enumeration, and the rest was immediately centrifuged (8,000 rpm for 5 min) and frozen for cytokine analysis. Next, the microtissue models were washed twice with PBS and lysed with 1% Triton X-100 (ThermoFisher) for 15 min at 37°C, followed by a manual physical disruption to CFU-enumerate the intracellular and attached/biofilm-like communities. As positive control, infected models without treatment were used, and uninfected samples were used as negative controls. Two microtissues per condition were used and three independent experiments were performed.

### Phage recovery assay

After the microtissue model was homogenised for CFU enumeration, the samples were centrifuged at 4,000 x g for 10 min for a phage counting spot assay on double layer agar. Briefly, three *E. coli* phage hosts (EA2, B31, and MH10) were grown overnight and cultured to log phase (OD_600_ = 0.15 – 0.2). Double-layer agar was then prepared by adding 100 μL of bacterial culture to 8 mL of 0.7% (w v^-1^) LB agar, then poured onto square LB 1.5% (w v^-1^) agar plates. The sample supernatant was serially diluted with SM Buffer [100 mM NaCl (Sigma-Aldrich, United Kingdom)], 8 mmMgSO4•7H2O (Sigma-Aldrich), 50 mM Tris-HCl pH 7.5 (Sigma-Aldrich), then spotted onto dried double layer-agar and incubated 37 °C overnight (∼18 h).

### LCPR1 planktonic killing assay (PKA)

*E. coli* UTI89 overnight cultures were diluted 1:100 either in MHB or 70% pHU:30% MHB (70% pHU:MHB) and added to flat-bottomed 96 well plates (Fisher Scientific, UK) in triplicate per treatment. The bacteria were grown to OD_600_ 0.15 (1 x 10^8^ CFU mL^-1^) and 100 μL of phage cocktail (containing 10^8^ PFU mL^-1^ of each individual phage) was added to each well, with dilution of phages made using MHB or 70% pHU:MHB. The PKA was performed at a multiplicity of infection (MOI) of 0.1, 1, and 10. The experiments were performed using the BMG Labtech SPECTROstar Omega, with measurements every 5 min (OD_600_) for 24 h, with shaking of 10 s before each reading. Three independent biological experiments were performed.

### Plug-Flow Linked Organoid (P-FLO) system

The P-FLO system utilised in this work was designed and manufactured in-house. A bespoke polydimethylsiloxane (PDMS) plug was designed for integration with 12-well plate Transwells (Corning) to allow continuous flow in the apical compartment of the insert. The Transwell plugs were manufactured by replica moulding using 3D-printed resin moulds. The material used for the plugs was PDMS (Sylgard^TM^ 184) at a 10:1 weight ratio of PDMS-to-curing agent. The computer-aided design (CAD) files for the moulds (in STL format) are freely available for download and use (available upon request). The mould model designs were sliced using PreForm software (Formlabs) and then 3D-printed using a Formlabs Form 3B+ SLA 3D printer in Grey resin V4 material (Formlabs). The print resolution was set to 50 μm. To enable effective curing and detachment of PDMS from the moulds, these were treated for 10 h at 80 °C and lined with a thin layer of silicon spray (HD-Sil Release, Ambersil,) before pouring of liquid PDMS. Once poured over the moulds, PDMS was allowed to cure overnight at 60 °C and the solidified part was subsequently removed from the moulds. A New Era NE-1000 syringe pump (New Era Pump Systems Inc., Farmingdale, NY, USA) and PTFE 16 mm diameter tubing (Darwin Microfluidics) were utilised to deliver media through the P-FLO system at the desired flow rate. 20 or 60 mL capacity syringes (Fisher Scientific) were used as a media reservoir in this work.

### Numerical simulation of the fluid dynamic field within the P-FLOW system

The P-FLO system was developed to simulate the levels of wall shear stress to which the bladder urothelium is subjected during bladder filling and voiding. Data on such wall shear stress values are underreported in the literature, due to impracticality of measurement and significant individual variability. A generally accepted estimation for shear stress on the bladder wall during filling is 0.02 N m^-2^. During voiding, the estimated wall shear stress increases 10- to 100-fold for a short period of time^31,32^. Two different volumetric flow rate values were thus selected for this work, to simulate bladder filling and voiding respectively. These were estimated by manual calculation to be 0.002 mL min^-1^ and 1.7 mL min^-1^, respectively. Subsequently, a CFD simulation was run on the P-FLO model to confirm the expected shear stress profile acting over the microtissue surface area. Given the relatively low flow rate necessary to obtain wall shear stress values in the 0.017 to 17 Pa range, the flow regime was modelled as laminar (the highest Reynolds number expected in the system is estimated to be ∼12).

A steady-state laminar flow simulation was carried out using COMSOL Multiphysics (COMSOL inc., Burlington, MA, USA) with the Laminar Flow (spf) physics interface. The model was set up to represent flow of water (modelled as incompressible and Newtonian fluid) through the P-FLO system. COMSOL solves the governing equations for laminar flow, consisting of the Navier–Stokes momentum equation and the continuity equation, expressed as:

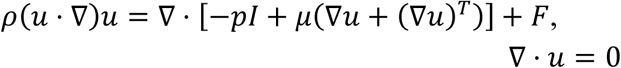

where *ρ* is the fluid density, ***u*** is the fluid velocity vector, *p* is the fluid static pressure, *μ* is the fluid dynamic viscosity, and ***F*** represents external body forces such as gravity. The equations were solved under steady-state conditions using the finite element method (FEM). Appropriate boundary conditions, including specified inlet mass flow rate values, no-slip wall condition, boundary layers, and outlet atmospheric pressure, were applied. The dependency of the numerical solution on the mesh element size was also evaluated, to identify the most appropriate meshing scheme.

### UTI89 infection under flow in the bladder microtissue model for 6h

Fully differentiated bladder microtissue models from day 16 were used for the infection. The bacterial inoculum was prepared as stated above. Then, the microtissues models were infected with *E. coli* UTI89-GFP (5.0 x 10^6^ bacterial cells mL^-1^), and the PDMS plugs were inserted into each Transwell, followed by the placement of a custom lid over the 12-well plate, created to feature 2 mm holes to fit the inlet and outlet PTFE tubing (Darwin Microfluidics). 20 mL syringes loaded with pooled urine were connected to the plugs’ inlets and placed into the multi-channel syringe pump. The outlets were connected to a waste reservoir, and the full system was placed into an incubator at 37°C with 5% CO_2_. A low flow rate was used to mimic the filling phase (0.02 mL min^-1^) for 6h, interrupted every 2 h with higher flow rate (0.7 mL min^-1^) for 1 min to mimic the voiding phase i.e. micturition. At the end of the experiment, the plugs were carefully removed, and the apical fluid was collected for CFU enumeration. Next, two washes with PBS were performed and microtissues were either lysed for CFU enumeration or fixed for imaging (see protocols below). Two microtissues per condition were used and three independent experiments were performed.

### Antimicrobial efficacy in the 3D-UHU under flow

Fully differentiated 3D-UHU models from day 16 were inoculated with *E. coli* UTI89-GFP (5.0 x 10^6^ bacterial cells mL^-1^), and, immediately after, the P-FLO system was connected. Infection was allowed to take place for 12h at 37°C with 5% CO_2_ under a flow rate (0.02 mL min^-1^) mimicking a filling cycle. After 12h, syringes were replaced with others containing pooled human urine either untreated, or containing antibiotics (nitrofurantoin, 256 μg mL^-1^), phage cocktail (LCPR1, 1 x 10^9^ phage mL^-1^) or the combination of both treatments (256 μg mL^-1^ and 1 x 10^9^ phage mL^-1^). During treatment, a low flow rate was used to mimic the filling phase (0.02 mL min^-1^) for 8h, interrupted every 2 h with a voiding cycle at the higher flow rate (0.7 mL min^-1^) for 1 min in each cycle. After plug removal, the fluid in the apical side was collected for CFU enumeration; the microtissues were washed twice with PBS, then lysed or fixed for imaging. Two microtissues per condition were used and three independent experiments were performed.

### Lactate Dehydrogenase (LDH) cytotoxicity assay

Cytotoxicity in the bladder microtissue model was assessed by quantification of the amount of lactate dehydrogenase (LDH) released into the apical compartment, using the commercially available CyQUANT^TM^ LDH Cytotoxicity Assay (Thermo Fisher). Calculation of percentage cytotoxicity was performed as recommended by the manufacturer. Models fully lysed with 1% Triton X-100 were used as a maximum LDH control. Fluorescence intensity was measured in a Tecan Spark® 10 M plate reader (λ_ex_ = 490 nm and λ_em_ = 680 nm). Three independent experiments were performed, including three technical replicates per experiment.

### Transepithelial electrical resistance

Transepithelial electrical resistance (TEER) of 3D-UHU was measured using the EVOM3 with a STX2-Plus electrode (World Precision Instruments, UK). Measurements were taken according to the manufacturer’s instructions. A custom 3D-printed holder was used to ensure the electrode was placed correctly in the Transwell insert (the holder design is available upon request). Blank readings were taken from an empty 12 mm, 0.4 μm pore polycarbonate membrane plastic insert (Corning) with 900 μL of CnT Prime Epithelial 3D Airlift Medium (CELLnTEC) in the apical compartment and 1.5 mL in the basolateral compartment. Post infection, 3D-UHU models were washed twice with PBS, and pre-warmed CnT Prime Epithelial 3D Airlift Medium was added in both apical and basolateral compartments (900 μL and 1.5 mL, respectively) before TEER measurements were taken. Three biological replicates were performed.

### Immunofluorescence staining and microscopy

Samples were fixed in 4% methanol-free formaldehyde (Fisher Scientific) in PBS and incubated at 4°C overnight. Then, the fixative solution was aspirated out, replaced with PBS, and stored at 4°C until staining. Sample membranes were cut out of the Transwells with a scalpel and placed in a 24-well plate with 500 μL PBS. Samples were then permeabilised with 0.2% Triton X-100 away from light for 35 mins at room temperature with rocking at 15 rpm. Samples were washed twice with PBS and blocked with 5% normal goat serum (NGS, Thermo Fisher) and 1% bovine serum albumin (BSA, Merck) in PBS for 1 h at room temperature with rocking at 15 rpm, washed once with 1% BSA, and primary antibodies were then added to each sample. Primary antibodies used were rabbit anti-cytokeratin 20 (PA5-22125, Invitrogen UK) diluted at 1:75 (11.73 μg mL^-1^) and mouse anti-uroplakin 3A (sc-166808, Santa Cruz) diluted at 1:50 (4 μg mL^-1^), both resuspended in 1% NGS with 1% BSA. Samples were incubated statically overnight at 4°C. Then, the samples were washed three times with 1% BSA. Secondary antibodies were then added: goat anti-mouse 488 (Invitrogen) diluted to 1:300 (6.67 μg mL^-1^) and goat anti-rabbit 555 (Invitrogen) diluted to 1:200 (10 μg mL^-1^) in 1% BSA, respectively, and incubated at room temperature for 2 h with rocking at 15 rpm. Samples were washed three times with PBS and incubated with phalloidin-647 (Invitrogen) diluted at 1:500 (from stock solution of 1 unit μL^-1^) in PBS for 3 h at room temperature with rocking at 15 rpm. 4’,6-diamidino-2-phenylindole (DAPI, Invitrogen) was added to the samples with phalloidin at 1:500 (2 μg mL^-1^) dilution and incubated in the same conditions for 30 min. Samples were washed 4 times with PBS and mounted onto glass microscope slides using ProLong™ Glass Antifade mounting media and covered with a cover slip. Slides were stored in a dark, dry place before imaging. Image visualization was performed by confocal laser scanning microscopy using an inverted microscope (Eclipse Ti, Nikon Inc, USA), equipped with a Digital Sight 10 microscope camera (Nikon) and X-cite LED for fluorescence imaging, using the 40x air-magnification lens and ×60 1.4 NA oil-immersion magnification lens. The microscope was controlled by NIS-Elements v.3.30.02 software.

To visualise intracellular invasion by confocal microscopy, an *E. coli* FITC-conjugated antibody (PA1-73029, Invitrogen) was used to detect only extracellular bacteria following our protocol previously described^22^. Briefly, the microtissues were incubated with anti-*E. coli* FITC-conjugated antibody at 1:50 in PBS (100 μg mL^-1^) for 6h at room temperature, rocking at 15 rpm. Following incubation, samples were washed 3 times with PBS after which the permeabilization and staining (phalloidin and DAPI) protocols (as described above) were carried out. Non-fluorescent bacteria (*E. coli* UTI89) or the RFP-tagged strain were used for these infections. Under confocal microscopy, the samples with FITC/RFP/DAPI signals corresponded to the extracellular communities, while the bacteria with only RFP/DAPI signals represented intracellular pathogens.

### IL-1β profiles

ELISA on supernatants was used to quantify the amount of IL-1β produced by 3D-UHU microtissues before and after infection (R&D Systems), according to the manufacturer’s instructions. Frozen supernatants were first diluted 1:10 with sterile PBS, and absorbance was measured in a Tecan Spark® 10 M plate reader at 450 nm. Three biological experiments were performed.

### Scanning electron microscopy

Samples were fixed in 2.5% glutaraldehyde in 0.1 M phosphate buffer (for 30 min at room temperature). They were then washed twice with 0.1 M of phosphate buffer, cut out of Transwells with a scalpel, and placed in a 24-well plate. Membranes were then dehydrated in a graded ethanol (Sigma-Aldrich) series: 10 min of incubation each in 30, 50, 70, and 90% and 10 min in 100% ethanol, twice. Dehydrated microtissues were completely dried using hexamethyldisilazane (HMDS, ThermoFisher). The membranes were detached from the plastic inserts using a scalpel, mounted onto aluminium stubs using conductive carbon tape, and sputter-coated with 7 nm of gold. SEM secondary electron (SE2) images were acquired at 2 kV with a Zeiss Sigma 300 Field Emission Gun Scanning Electron Microscope (FEG-SEM) using Zeiss SmartSEM v7.02. False colouring of the images was performed using GIMP 2.10 software.

### Bacterial cell elongation quantification

Confocal *z*-stacks of the GFP channel (*E. coli* UTI89-GFP) were acquired from the apical surface of the urothelium using a Nikon Ti2 confocal microscope with a 60× objective after 6h under flow or static fluidic conditions. To analyse bacterial morphology, 3D images were processed in Ilastik^33^ using a supervised machine-learning classifier to generate segmentation masks, with the model trained to exclude poorly segmented regions and objects not corresponding to single bacteria. The resulting 3D masks were imported into FIJI^34^ and analysed with the MorphoLibJ^35^ plugin to extract morphological parameters, including geodesic diameter (i.e., a measure of cell length), sphericity, surface area, and volume. Three biological replicates (6 images per biological replicate) were analysed independently with at least 5,000 bacterial objects segmented per condition.

### Image analysis of bacterial coverage

Maximum-intensity projections of the GFP channel, corresponding to *E. coli* UTI89-GFP, were generated from confocal *z*-stacks acquired using a Nikon Ti2 confocal microscope with a 40× objective. Segmentation of bacteria-containing regions was performed using the Labkit^36^ plugin (FIJI, built-in) with manually trained pixel classifiers to generate binary masks. The resulting masks were binarized in FIJI, and the bacterial coverage area was quantified as the proportion of GFP-positive pixels relative to the total image area. Six images were analysed per biological replicate (*n* = 3).

### Statistical analysis

Data were analysed and plotted using GraphPad Prism version 10 and RStudio 2025.09.1+401. The statical test performed for each data set are indicated in the corresponding figure legends.

## Results

### Antimicrobial susceptibility testing (AST) against UPEC strains in caMHB vs human urine

Antibiotic efficacy is still primarily assessed using a single *in vitro* bioassay in standard growth medium^37^. However, this assay overlooks key environmental factors present during host-pathogen interactions *in vivo* that profoundly influence drug performance^38^. We investigated the AST profiles of clinically relevant antibiotics against several *E. coli* strains including clinical UPEC isolates (EC3, EC4, ClinA), asymptomatic strains (HM50, COM1, COM2), and reference strains such as the commonly studied pyelonephritis strain (CFT073) and cystitis strain (UTI89), alongside a non-pathogenic lab strain (MG1665). The selected antibiotics are commonly used to treat both complicated and uncomplicated UTIs following UK and European clinical guidelines^39^. In addition to the gold-standard growth media (caMHB), we included 70% pooled human urine (pHU) (in 30% caMHB) to mimic a UTI infection microenvironment as previously reported^40^. Prior to AST evaluation, bacterial growth was evaluated in caMHB, 70% pHU and 100% pHU (**Figure 1A**). While growth in 100% urine was markedly reduced, cultures in 70% pHU exhibited similar growth kinetics to caMHB, supporting the use of 70% pHU as a physiologically relevant medium for subsequent antimicrobial testing.

**Figure 1.**
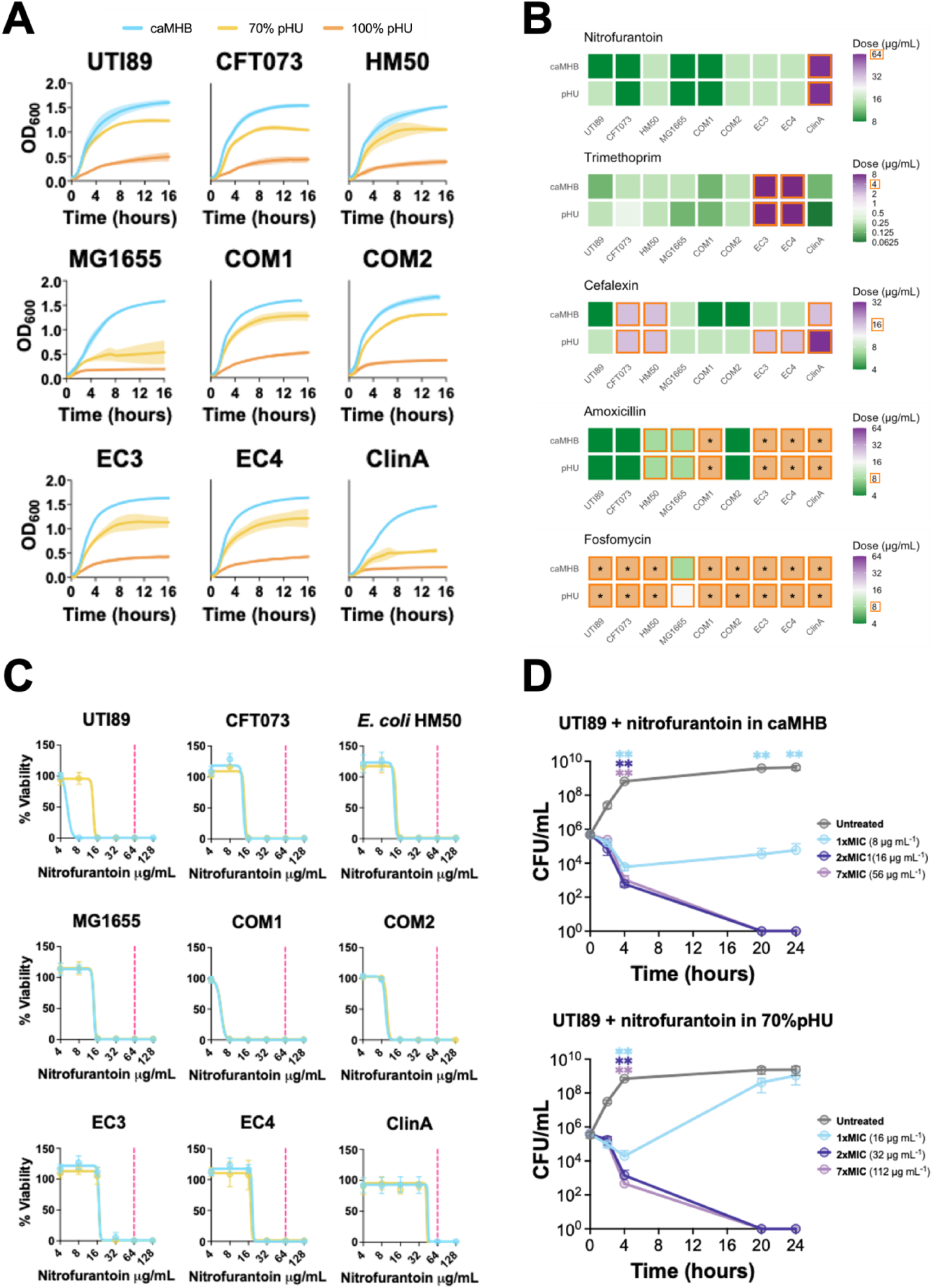
**(A)** Growth curves of *E. coli* strains in caMHB (blue), 70% pHU (yellow) and 100% pHU (orange). The shadowed colour represents the standard deviation between the three biological replicates, and the line represents the mean. **(B)** Heat map of the MIC results of nitrofurantoin, cefalexin, amoxicillin and fosfomycin against *E. coli* strains in caMHB and 70% pHU. The orange square box defines when the MIC value sat on the EUCAST breakpoint or above, and the * means the MIC could not be determined because bacterial growth was observed at the highest concentration tested (above the EUCAST breakpoints). **(C)** Dose response graphs of *E. coli* strains in caMHB (blue) and 70% pHU (yellow) against nitrofurantoin. The dashed pink line defines the EUCAST breakpoint. **(D)** Time killing assay of *E. coli* UTI89 against nitrofurantoin in caMHB (top). A t-test was performed to compare the CFU of the individual treatments with the CFU of the untreated controls. Time killing assay of *E. coli* UTI89 against nitrofurantoin in 70% pHU (bottom). A t-test was performed as above. For graphs, the error bar represents the standard deviation between the three biological replicates, and the empty dots represent the mean.

The MIC results are summarised in **Figure 1B**. All strains showed similar levels of susceptibility to nitrofurantoin (8-16 μg mL^-1^), except ClinA with an MIC of 64 μg mL^-1^ (at the breakpoint defined by the The European Committee on Antimicrobial Susceptibility Testing [EUCAST]). Trimethoprim was active against all strains except EC3 and EC4, where the MICs were above the breakpoint in both caMHB and 70% pHU. In the case of fosfomycin, amoxicillin and cefalexin, 5 of the 8 strains exhibited MICs above the resistance breakpoints. In general, similar trends for all antibiotics tested were observed in caMHB and 70% pHU, with non-significant differences between the two microenvironments. In constrast, for cephalexin, EC3, EC4 and ClinA showed higher MICs in pHU in comparison with caMHB, crossing the susceptibility breakpoint when pHU was used. These results suggest that the presence of urine can affect AST depending on the strain and antibiotic tested, but that these effects are not drastic in an *in vitro* environment.

Nitrofurantoin and trimethoprim are the most common antibiotics prescribed in the UK to treat uncomplicated UTIs,^41^ and they exhibited the greatest antimicrobial performance against the *E. coli* strains tested in this study. Nitrofurantoin remains a resilient therapeutic option for UTIs, due to its localized delivery, favourable pharmacokinetic and pharmacodynamic properties, and multifaceted effects on bacterial physiology^42^. Resistance acquisition is constrained by the need for multiple independent mutations and the near-absence of horizontally transferable resistance mechanisms^43^. Given that there is higher incidence of *E. coli* resistance to trimethoprim compared with nitrofurantoin^44^, making the latter more clinically applicable, we decided to focus on nitrofurantoin in the subsequent steps of the study

Next, we determined the minimum bactericidal concentration (MBC) by measuring bacterial viability as shown in **Figure 1C** and **Figure Supplementary 1**. The MBC values for nitrofurantoin were identical between caMHB and 70% pHU for all the strains except UTI89, with 1-fold increase in the urine microenvironment. Hence, we further investigated the killing kinetics of nitrofurantoin against *E. coli* UTI89, comparing caMHB and 70% pHU microenvironments (**Figure 1D**). The killing kinetics showed distinct media-specific profiles, especially at 16 μg mL^-1^. In caMHB, nitrofurantoin at MIC, 2xMIC and 7xMIC for 4h caused a statistically significant CFU reduction in comparison with the untreated control (all *p* < 0.001**). At 20h and 24h, nitrofurantoin at MIC reduced significantly CFUs in comparison with the untreated control (all *p* < 0.001**), while 2xMIC and 7xMIC caused total bacterial eradication and statistical tests were not performed. In the pHU, nitrofurantoin at MIC, 2xMIC and 7xMIC at 4h caused a statistically significant CFU reduction in comparison with the untreated control (all *p* < 0.001**). At 20h and 24h, nitrofurantoin at MIC was not significantly different to the untreated control while 2xMIC and 7xMIC caused total bacterial eradication. In both microenvironments, at least 2xMIC was needed to eradicate the bacterial viability. However, in the case of 70% pHU, a higher dose of nitrofurantoin was necessary to achieve bacterial eradication (4-fold increase in drug concentration) in comparison with caMHB. These results suggest a modest effect of the urinary microenvironment on nitrofurantoin response in the *in vitro* conditions evaluated.

### Effect of nitrofurantoin against *E. coli* UTI89 in a human bladder microtissue model under static conditions

To better understand treatment failure in UTI, we used our urine-tolerant human urothelial (3D-UHU) microtissue model, which recapitulates key features of the host microenvironment and host-pathogen interactions, as a novel platform for antimicrobial treatment screening^7,22^. Following the experimental setup outlined in **Figure 2A**, the urothelium was infected with *E. coli* UTI89, which exhibited intracellular invasion events at 20 h post-infection (hpi) as shown in **Figure Supplementary 2A-D**, consistent with our previous observations.^22^ Infected tissues were then treated with two consecutive doses of antimicrobial agents in the apical urine compartment (Tx1 and Tx2), each administered for 4 h. After take-down, we assessed both the apical (urine, planktonic bacteria) compartment and the microtissue (adhered and intracellular bacteria) compartment.

**Figure 2.**
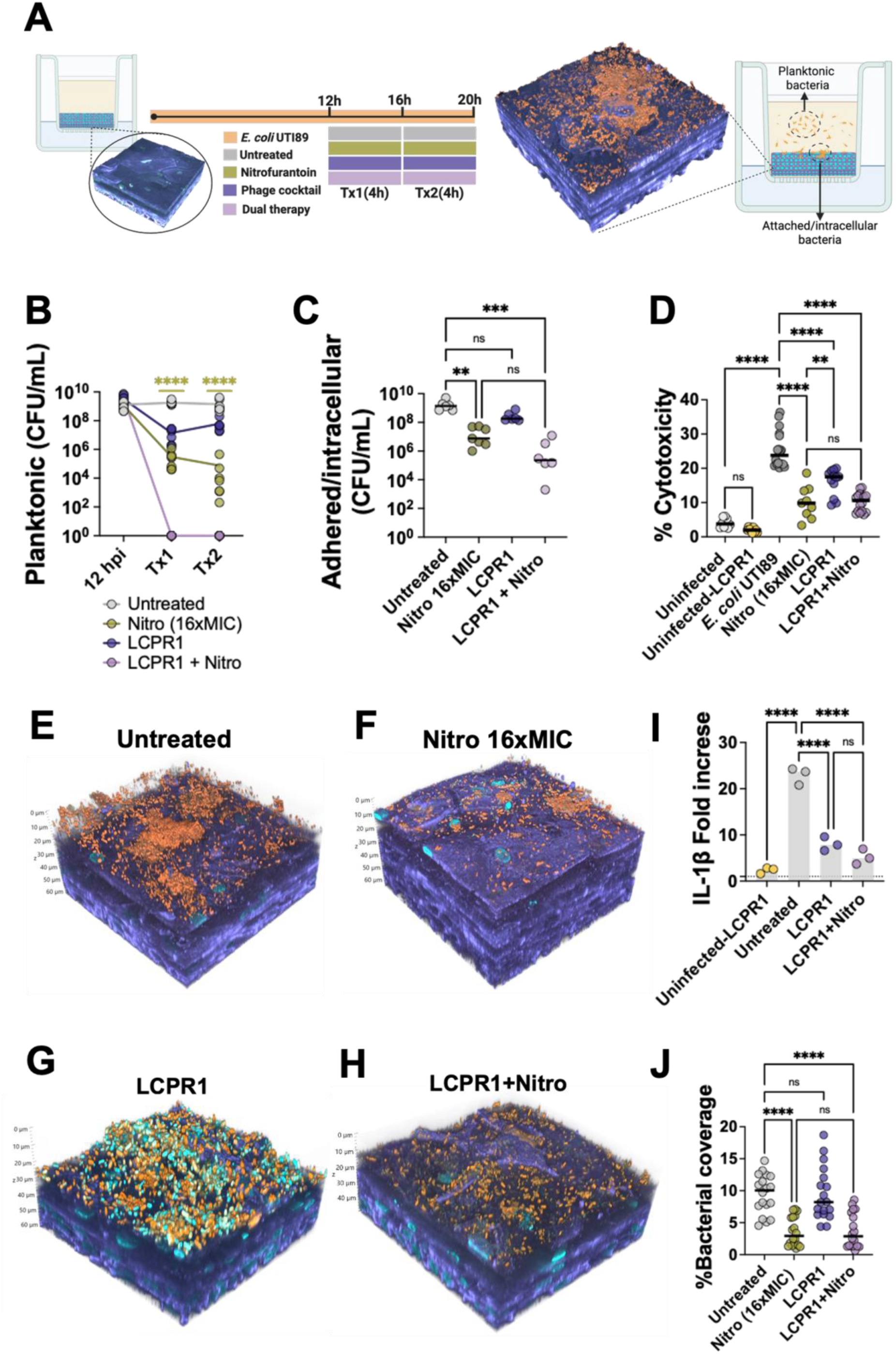
**(A)** Diagram of the 3D-UHU microtissue model infected with *E. coli* UTI89 followed by antimicrobial treatment. Fully differentiated organoids (d≥16) were infected with *E. coli* UTI89-GFP at 5.0 x 10^6^ bacterial cells mL^-1^ for 14h. Then, the apical compartment was changed every 4 h with pooled human urine either untreated (as shown in grey) or supplemented with nitrofurantoin (256 μg mL^-1^ in green), LCPR1 phage cocktail (blue), and the combination of both treatments (purple). Post treatments, the planktonic bacteria in the apical compartment and the attached or intracellular bacterial populations were investigated as described in the drawing. **(B)** Bacterial load by CFU enumeration in the apical side (planktonic) after 12h infection, post-4h 1^st^ treatment (Tx1) and post-4h 2^nd^ treatment (Tx2). A Kruskal-Wallis test was performed after Tx1 to compare nitrofurantoin, LCPR1 and combinatorial treatment with the untreated control. **(C)** Bacterial load by CFU enumeration of the attached/intracellular bacteria after the two consecutive treatments. A Kruskal-Wallis test was performed as above. **(D)** Cytotoxicity caused by *E. coli* UTI89 under antimicrobial treatment was assessed by lactate dehydrogenase (LDH) release assay at 20 hpi. Data are plotted as means (line) and the empty dots represent the technical replicates from three biological experiments. An ordinary Anova test was performed to compare the different groups. **(E)** Representative 3D rendering of confocal microscope images of *E. coli* UTI89 untreated, exposed to nitrofurantoin **(F)**, LCPR1 **(G)** and dual treatment **(H)** in 3D-UHU. UTI89-GFP is represented in orange, the nucleic acid staining in cyan, and F-actin staining in purple. **(I)** Cytokine IL-1β production by the bladder microtissue at 20 hpi. Fold changes compared to uninfected controls are plotted as means ± standard deviation of the three biological triplicates. An ordinary Anova test was performed to compare the different groups. **(J)** Bacterial area coverage of the urothelium obtained from confocal imaging. Data plotted as means (line) and each empty dot represents a technical replicate from the three biological experiments. An ordinary Anova test was performed to compare the different groups.

Firstly, we assessed the efficacy of nitrofurantoin at a clinically relevant concentration^43^ (16xMIC, 256 μg mL^-1^). Treatment significantly reduced planktonic bacterial burden in the apical compartment, with a ∼3-4 log CFU reduction observed in both Tx1 and Tx2 compared with the untreated control (**Figure 2B**; p < 0.0001, ***). In contrast, nitrofurantoin treatment against urothelium-associated bacteria, which included attached and intracellular bacterial populations, was somewhat less effective, manifesting a ∼2-log reduction (**Figure 2C**; p =0.0059,**); this finding was consistent with reductions observed in the confocal imaging (**Figure 2E-F**) and the subsequent bacterial coverage area analysis (**Figure 2J**). Additionally, nitrofurantoin conferred a host-protective effect, as evidenced by reduced LDH release (p<0.0001***) relative to untreated controls (**Figure 2D**). Intracellular bacterial communities (IBCs) were observed after antibiotic treatment by confocal imaging, indicating that intracellular reservoirs can survive nitrofurantoin exposure, which might cause eventual treatment failure even if the bacterial burden was reduced in both planktonic and urothelium-associated bacteria (**Figure Supplementary 3**).

The antimicrobial effect of nitrofurantoin was dose dependent. At 7xMIC (112 μg mL^-1^), no significant difference in bacterial burden was observed in the attached/intracellular bacterial populations. In the planktonic population, a significant CFU reduction (p<0.001, ***) was observed after the first treatment exposure (Tx1) whilst bacterial re-growth was observed after the second exposure (Tx2) (**Figure Supplementary 2E-F**). Post-infection, the bacterial population exposed to the bladder microtissue environment (challenged with 7xMIC nitrofurantoin or untreated) was recovered and exposed to nitrofurantoin using a MIC microdilution method with standard growth medium. The MIC remained unchanged and both groups exhibited an identical MIC before and after the exposure to the bladder microenvironment (**Figure Supplementary 2G**), confirming that antimicrobial resistance mutations had not occurred during the course of the experiment. Despite this confirmed *in vitro* drug-sensitivity, treatment at concentrations above the MIC (7xMIC) failed to reduce the bacterial burden in the urothelial compartment, exemplifying the discrepancy between simplified *in vitro* tests and platforms that mimic the host microenvironment and host-pathogen interactions.

### Phage cocktail (LCPR1) and nitrofurantoin combinatorial therapy against *E. coli* UTI89 in a human bladder microtissue model under static conditions

Following evaluation of conventional antibiotic therapy, we next investigated the potential of bacteriophage-based therapy to target *E. coli* UTI89. The growth kinetics of UTI89 was monitored under the exposure of a bacteriophage cocktail (LCPR1) using different MOI in two different media. The bacterial growth was impaired up to 10 h for all MOIs tested in Müller-Hinton broth (MHB). After this period, partial regrowth was observed (OD_600_ ∼0.5), while the untreated control reached full growth (OD_600_ ∼2) (**Figure Supplementary 4A**). In contrast, LCPR1 efficacy was markedly reduced in 70% pHU:30% MHB, with no significant growth inhibition observed at MOI 1 and 10. Only at MOI 100 partial growth inhibition was observed (**Figure Supplementary 4B**). After 24h, the CFU reduction caused by LCPR1 was similar in both microenvironments (**Figure Supplementary 4D-E**) where only MOI=100 caused ∼2 log CFU reduction, as bacterial re-growth was also observed in the MHB after 12 h. Phage quantification from the killing assay demonstrated varying amplification levels after 24 h for each *E. coli* phage (Leic00, JK03, and UP17). Overall, phage recovery in MHB was lower than in 70% pHU for all three phages (**Figure Supplementary 4C,F**). This corresponded with greater reduction of bacterial populations in MHB in the first 10 h of phage exposure than in 70% pHU (**Figure Supplementary 4A**), suggesting that phage efficacy was greater in nutrient-rich media. A reduction of approximately 1–2 log units in phage titres was also observed in the uninfected control in 70% pHU. These findings underscore the influence of the type of culture medium on phage activity, suggesting that host-like conditions may limit therapeutic efficacy compared with standard laboratory media as previously reported by others^45,46^.

Next, we evaluated the effect of LCPR1 (1 x 10^9^ phage mL^-1^) against *E. coli* UTI89 using the static 3D-UHU microtissue infection model (**Figure 2A**). LCPR1 failed to significantly reduce bacterial burden (p=0.0971, ns) and colonization of the urothelium (p=0.8377, ns) in comparison with the untreated control (**Figure 2B-C,E,G,J**). Interestingly, a reduction in LDH release was observed in the LCPR1 treated group (p<0.0001***), indicating decreased cytotoxicity relative to the untreated control (**Figure 2D**). However, this protective effect was less pronounced than that conferred by nitrofurantoin (p=0.0043**). Notably, the reduction in LDH release did not correlate with bacterial load, suggesting a mechanism independent of direct bactericidal activity. To explore potential host-modulatory effects, we quantified levels of the pro-inflammatory cytokine IL-1β by ELISA. LCPR1 treatment resulted in a statistically significant decrease in IL-1β secretion (p<0.0001***) compared with the untreated group (**Figure 2I**), supporting the hypothesis that phage exposure may attenuate host inflammatory responses. Together, these findings suggest that while LCPR1 lacks bactericidal efficacy in this model, it may exert beneficial immunomodulatory effects that reduce tissue damage, although perhaps at the expense of antimicrobial effects.

Given the antimicrobial activity of nitrofurantoin and the host-modulatory effects observed with LCPR1, we next assessed the impact of combining both treatments to determine whether dual therapy could enhance bacterial clearance and mitigate host tissue damage. The dual treatment caused a total bacterial eradication in the apical compartment, and it achieved a ∼4-log CFU reduction (p<0.0001***) in the microtissue compartment in comparison with the untreated control, which is considered clinically relevant^47^ (**Figure 2B-C,E-H,J**). The dual therapy was, however, not superior to the antibiotic treatment (p>0.9999, ns) in terms of host-protection (cytotoxicity). In the aspect of immunomodulation the dual treatment was not superior (p=0.2625, ns) to the LCPR1 treatment (**Figure 2D,I**).

Phage recovery from the 3D-UHU was also assessed in both the apical (Tx1 and Tx2) and in the microtissue compartments (**Figure Supplementary 4 G-I**). Comparable titres were observed in the apical compartments (Tx1 and Tx2) for phage Leic001 and UP17 for single LCPR1 with combined treatment (**Figure Supplementary 4 G,H**). In contrast, JK03 showed higher titres in the LCPR1 single treatment compared with the combined treatment. In the microtissue compartment all three phages recovered from LCPR1 alone had higher titres compared to the dual treatment (**Figure Supplementary 4I)**. This result correlated with the bacterial count result, which showed that a combined treatment effectively reduced bacterial population compared to LCPR1 alone (**Figure 2B-C**), and subsequently reduced the amplification level of phage. Nevertheless, this pattern was not observed in the apical phage recovery results, which highlights the complexity of phage-bacteria dynamics within the bladder microenvironment.

### Fluid flow triggers UPEC elongation in a human bladder microtissue model

To recapitulate the mechanical forces generated by urine flow, we engineered a mesofluidic platform referred to as the Plug-Flow Linked Organoid (P-FLO) system (**Figure Supplementary 5A**), designed to introduce physiologically relevant levels and temporal profiles of wall shear stress into the 3D-UHU microtissue model^22,23^. The system enables cyclic changes of flow rate (and hence of applied wall shear stress) in the apical surface of the differentiated urothelial microtissue, mimicking the filling and voiding phases of the micturition cycle (**Figure Supplementary 5A**). Notably, application of flow (voiding-filling cycles for 6h) to uninfected 3D-UHU models did not alter barrier integrity or expression of the key differentiation markers CK20 and UPIII, compared with static controls (**Figure Supplementary 5C–E**).

To investigate the impact of fluid dynamics on UPEC infection, 3D-UHU models were infected with *E. coli* UTI89-GFP and, immediately following infection, a continuous flow of 0.002 mL min^-1^ was applied for 6 h to simulate bladder filling. To mimic urination cycles, this flow was interrupted every 2 h by a 1-min voiding event, at an increased flow rate of 0.7 mL min^-1^ (**Figure 3A**). This novel experimental setup recapitulates key mechanical cues of the bladder environment, enabling assessment of infection progression under conditions that approximate *in vivo* wall shear stress levels as well as the dynamic nature (i.e. alternating low and high wall shear stress) of the micturition cycle. Numerical simulations were performed to investigate the shear stress field generated on the urothelium at the corresponding volumetric flow rates tested. The computational modelling results revealed that the area of highest shear stress was near the inlet during the voiding phase, with intermediate shear stress levels near the outlet region. During the filling phase, the wall shear stress field was more uniformly distributed and exhibited the lowest values among all simulated conditions (**Figure Supplementary 6A-B**). After the filling-voiding cycles (6 hpi), image analysis revealed comparable levels of bacterial attachment in regions that were exposed to high (HSS) and intermediate (ISS) wall shear stress levels during the filling phase (**Figure Supplementary 6D,F**). Therefore, image analyses were subsequently conducted on randomly selected areas within the microtissue. In addition, a flow protocol that did not include voiding phase was evaluated, to assess whether the latter had a major impact on bacterial cell morphology and behaviour (**Figure Supplementary 6A**). It was found that the filling phase alone (6 hpi), associated with regions of lowest shear stress (LSS) values, caused increased bacterial adhesion in comparison with samples exposed to filling-voiding cycles (**Figure Supplementary 6D,F**).

**Figure 3.**
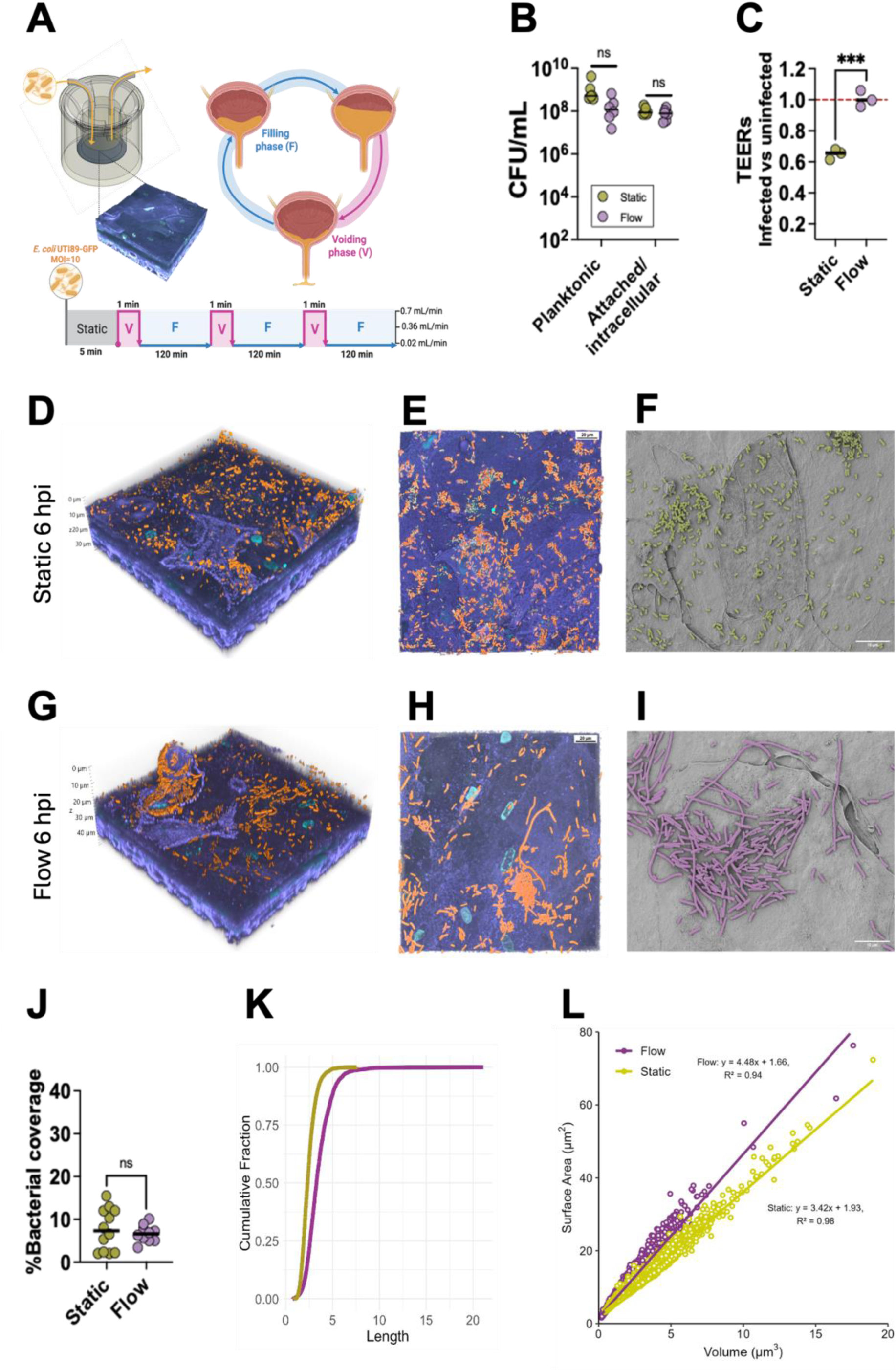
**(A)** Schematic diagram of the P-FLO system integrated with the Transwell-grown 3D-UHU bladder microtissue model, programmed to deliver controlled volumetric amounts of human urine over defined time periods to mimic the filling and voiding phases of urination. Fully differentiated models (d≥16) were infected with *E. coli* UTI89-GFP at 5.0 x 10^6^ bacterial cells mL^-1^ and, immediately after, the P-FLO system was used to incorporate flow dynamics for 6h with a 0.002 mL min^-1^ flow rate (mimicking filling), which was interrupted every 2h by a voiding phase of 1 min with a greater flow rate of 0.7 mL min^-1^. **(B)** CFU enumeration of *E. coli* UT89-GFP on the apical side (planktonic) and attached/intracellular population associated with the bladder urothelium (6 hpi) in static conditions and under flow. The black line represents the mean and the dots the technical replicates. Mann-Whitney tests were performed to compare the static vs flow groups in the apical and microtissue-associated bacteria, with no significant difference in both cases (p values of 0.53 and 0.45, respectively). **(C)** Epithelial barrier integrity measured by TEER (6 hpi) comparing static and flow-dynamic infection conditions. The data is presented as a ratio between the uninfected control and the infected samples obtained in static or flow conditions. A ratio equal to 1 (shown as a red dashed line) indicates that the epithelial barrier integrity is uncompromised and values <1 correspond to compromised barrier. **(D)** Representative 3D rendering and **(E)** top-view confocal microscopy images of *E. coli* UTI89-GFP 6 hpi under static and flow conditions **(G-H)**. UTI89-GFP is represented in orange, the nucleic acid staining in cyan, and F-actin in purple. **(F)** False-coloured SEM micrograph of *E. coli* UTI89-GFP 6 hpi in static and **(I)** fluid dynamic conditions. **(J)** Bacterial area coverage of the urothelium obtained from confocal imaging under static and flow dynamic conditions. **(K)** Cumulative fraction plot of the bacterial length obtained under static vs dynamic fluidic conditions, obtained from the 3D bacterial cell segmentation image analysis. **(L)** Surface area vs volume plot obtained from the 3D bacterial cell segmentation image analysis.

Quantification of CFUs revealed no significant differences in planktonic bacteria within the apical compartment or in urothelium-associated bacterial populations at 6 hpi under filling-voiding cycles versus static conditions (**Figure 3B**). Despite comparable bacterial loads, barrier integrity was significantly better preserved under flow, suggesting that mechanical cues mitigate infection-induced epithelial disruption (**Figure 3C**). Bacterial attachment to the urothelium was not affected after the voiding-filling cycles in comparison with the static control (**Figure 3J**). However, *E. coli* UTI89 underwent marked bacterial elongation/filamentation, as observed by both confocal and electron microscopy under the filling-voiding cycles 6 hpi (**Figure 3D-I**). To assess the impact of flow dynamics on bacterial morphology during infection, the cumulative fraction of bacterial objects as a function of their length under static and dynamic (flow) conditions was quantified using 3D-bacterial segmentation image analysis. The cumulative distribution curves revealed distinct bacterial size distribution profiles between the two conditions (**Figure 3K**). Under static conditions, bacterial objects rapidly reached full representation, with the cumulative fraction approaching 1.0 at shorter lengths (mustard-coloured curve). In contrast, flow-exposed bacteria exhibited a broader size distribution, with a more gradual accumulation of longer objects (purple-coloured curve), indicating delayed saturation of the cumulative fraction. Additionally, UPEC under flow exhibited a higher surface area-to-volume ratio further indicating greater elongation than those cultured statically (**Figure 3L**). These findings confirm that flow promoted elongation of bacterial cells, consistent with enhanced filamentation or altered growth dynamics in response to mechanical cues. In the HSS and ISS regions after voiding-filling cycles, the extent of bacterial cell elongation was comparable (**Figure Supplementary 6E,F**). To assess whether the observed bacterial elongation was a consequence of the alternating filling-voiding cycles (and the associated temporal change in wall shear stress levels), we tested a flow protocol without the voiding phases (**Figure Supplementary 6E,F**). The filling phase alone (6 hpi) induced cell elongation as observed in the confocal image (**Figure Supplementary 6C,F),** suggesting that the low wall shear stress regimen representative of bladder filling was sufficient to induce bacterial elongation and that alternation of low and high wall shear stress levels was not the primary mechanism.

### Fluid flow affects antimicrobial susceptibility (for both bacteriophage and antibiotics) against UPEC in the 3D-UHU urothelial microtissue model

Given that more physiological flow dynamics allow continuous nutrient exchange and assert wall shear stress levels that could influence pharmacodynamics, we next used the P-FLO 3D-UHU system, to determine how flow affected the antimicrobial susceptibility of UPEC during infection. After the microtissue was infected with *E. coli* UTI89-GFP, a continuous flow of 0.002 mL min^-1^ was applied for 12 h to simulate bladder filling. Following the 12 h infection, either pHU only, or pHU containing nitrofurantoin (16xMIC, 256 mg mL^-1^), LCPR1 (1 x 10^9^ phage mL^-1^) and a dual treatment containing nitrofurantoin and LCPR1, was applied. To mimic urination cycles, the flow was interrupted with 1-minute voiding events at an increased flow rate of 0.7 mL min^-1^ every 2 h for a total of 8h (**Figure 4A**). The addition of flow in the 6 hpi resulted in bacterial cell elongation and a similar bacterial burden in the 3D-UHU in comparison with the static infection (**Figure 3D-J**). Similarly, at 20 hpi, the cell elongation phenomenon was maintained (**Figure 4B**) and greater bacterial attachment and burden (1-log CFU increase) were observed at 20 hpi in comparison with 6 hpi (**Figure 4D,I** and **Figure 3B).** Image analysis at 20 hpi (**Figure 4J**) demonstrated significantly greater bacterial attachment to the urothelium under flow conditions (22% bacterial coverage) than under static conditions (**Figure 2J**) with a 10% coverage. Wall shear stress promoted UPEC attachment compared with our previously established static infection model (in the absence of flow), indicating that hydrodynamic forces enhance adhesion and colonization in the 3D-UHU microtissue model (**Figure 4B,E,J, Figure 2E,J** and **Figure Supplementary 7D-E**). Furthermore, the bacterial invasion strategy of UPEC previously observed in static conditions (**Figure Supplementary 2**) was not impaired under flow conditions, as IBCs were observed under mechanical stress cues (**Figure 4C, Figure Supplementary 7A-C**).

**Figure 4.**
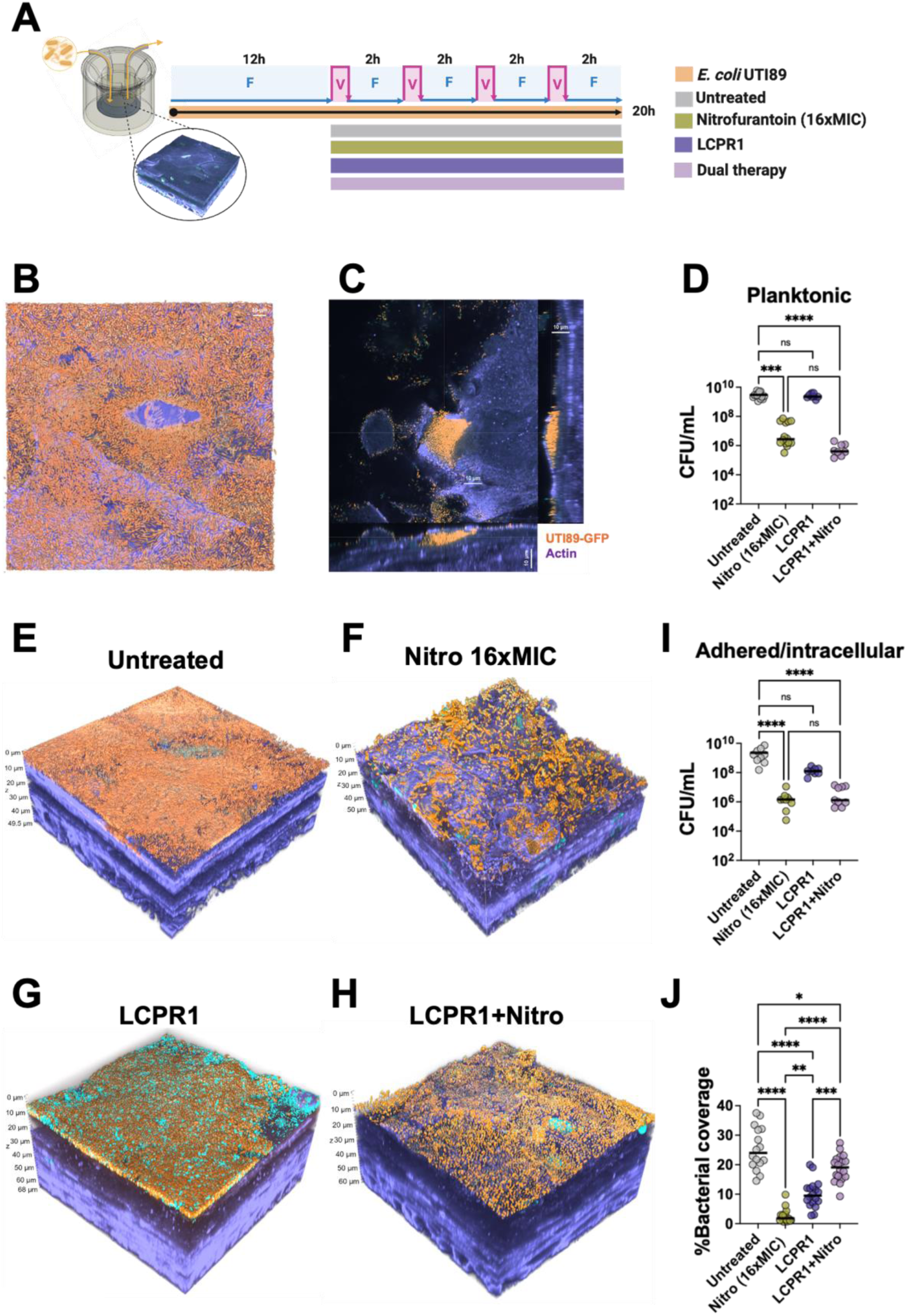
**(A)** Diagram of the 3D-UHU microtissue model infected with *E. coli* UTI89 followed by antimicrobial treatment under flow conditions using the P-FLO system. Fully differentiated organoids (d ≥16) were infected with *E. coli* UTI89-GFP at 5.0 x 10^6^ bacterial cells mL^-1^ for 12 h with a 0.002 mL min^-1^ flow rate mimicking filling. Then, the infected bladder microtissues were challenged with antimicrobial therapies. To mimic urination cycles, the flow was interrupted by 1-minute voiding events at an increased flow rate of 0.7 mL min^-1^ every 2 h, for a total of 8 h. **(B)** Top view of a representative 3D rendering of confocal microscopy image of *E. coli* UTI89-GFP untreated at 20 hpi under flow in the 3D-UHU. **(C)** Orthogonal view of an IBC under flow conditions. UTI89-GFP is represented in orange, the nucleic acid staining in cyan, and F-actin in purple. **(D)** Bacterial load by CFU enumeration in the apical side (planktonic) at 20 hpi. A Kruskal-Wallis test was performed to compare nitrofurantoin, LCPR1 and combinatorial treatment with the untreated control. **(E)** Representative 3D rendering of confocal microscope images of *E. coli* UTI89 untreated, exposed to nitrofurantoin **(F)**, LCPR1 **G**) and dual treatment **(H)** in 3D-UHU 20 hpi under flow conditions. **(I)** Bacterial load by CFU enumeration of the attached/intracellular bacteria post-treatment under flow conditions. **(J)** Percentage of bacterial area coverage on top of the urothelium obtained from confocal imaging. For all graphs, data are plotted as means (line) and each empty dot represents a technical replicate (different images) from the three biological experiments.

Nitrofurantoin significantly reduced planktonic bacterial burden in the apical compartment, with a ∼3-log CFU compared with the untreated control (**Figure 4D**; p < 0.0001,***). Against urothelium-associated bacteria, nitrofurantoin achieved a ∼3-log reduction (**Figure 4I**; p < 0.0001,****). Nitrofurantoin also exhibited distinct killing kinetics against planktonic and urothelium-associated uropathogenic populations under static versus flow conditions. In static conditions, a greater reduction in planktonic CFUs (p=0.0019, **). was observed, whereas under flow, nitrofurantoin demonstrated a non-significant antimicrobial activity (p=0.1547) against urothelium-associated bacteria in comparison with static conditions (**Figure Supplementary 7F**). Notably, despite increased overall drug exposure under flow, the antibiotic activity against planktonic bacteria was diminished.

LCPR1 failed to reduce bacterial burden under flow conditions to the same extent observed in static environments (**Figure 4D,I**). The dual treatment did not outperform the antimicrobial effect of nitrofurantoin alone, with comparable bacterial loads detected in both planktonic and urothelium-associated populations under flow (**Figure 4D,I**). Under static conditions, the CFU data (**Figure 2C**) were consistent with the bacterial coverage analysis (**Figure 2J**). In contrast, under flow conditions, the bacterial coverage (**Figure 4J**) did not align with CFU counts (**Figure 4I**), indicating a divergence between bacterial attachment and viability in the presence of shear stress.

The most striking difference in treatment efficacy emerged when comparing the dual therapy’s impact on planktonic bacteria between static and flow conditions (p<0.0001, ****). While complete eradication was achieved in static environments, flow conditions yielded only a ∼4-log reduction, highlighting how mechanical forces and microenvironmental cues under flow conditions might impact host-pathogen interactions and bacterial physiology, thereby modulating the overall antimicrobial efficacy (**Figure Supplementary 7F**). Compared with the static model, phage titres were similar overall, with lower phage counts in the adherent/intracellular compartment (**Figure Supplementary 7G-H**). Phage recovery of the planktonic compartment from the flow model showed higher titres in the LCPR1-only group than in the dual-treatment, consistent with the lower bacterial counts observed under dual treatment (**Figure Supplementary 7G, Figure 4D,I**). Notably, phage UP17 showed reduced titers levels both in the apical and microtissue compartment, which also reflects lower infectivity compared to the other two phages, JK03 and Leic001 (data not shown from independent experiments).

## Discussion

At least ∼260,000 deaths were associated with UTI linked to AMR cases in 2019^48^. Recurrent UTIs are extremely common, with ∼ 30% of all women experiencing a recurrence within one year of their original infection, and some of these experiencing repeated infections for years^49^. Poor treatment outcomes despite appropriate antibiotic therapy highlight the limited predictive value of standard *in vitro* antimicrobial susceptibility testing^14^. Heithoff *et al*., showed that, by recapitulating more closely the physiological conditions of infection (i.e. presence of urine, serum or cell culture media) in comparison with standard bacterial growth media (MHB), it was possible to achieve a closer prediction of antimicrobial performance in the clinical setting.^40^ In a similar manner, we showed that three UPEC isolates had an increased MIC against cefalexin, crossing the defined EUCAST breakpoint when 70% pHU was used. ^40^ Since we used the cationic-adjusted version of MHB (caMHB) with divalent cations at physiological concentrations, a less marked difference between 70% pHU and caMHB microenvironments was expected. Nitrofurantoin demonstrated the most potent antimicrobial activity against all the isolates tested in both microenvironments. However, when the complexity of the assay was increased (time-killing assays), the effect efficacy of nitrofurantoin against UPEC UTI89 was reduced in the urine microenvironment.

Advances in the bioengineering and microfluidic fields in the last decade have led to the development of organoids and organ-on-chip technologies which provide a physiologically relevant platform to investigate infectious diseases^50,51^. Recently, human airway microtissue models have revealed that *Pseudomonas aeruginosa* invades the epithelium via goblet cells^52^ and faces a trade-off between mucosal colonization and antibiotic tolerance^53^, highlighting the potential of organoid platforms as physiologically relevant systems for studying infection dynamics and testing antimicrobial therapies. Here, by employing our 3D-UHU microtissue model, we revealed that, whilst nitrofurantoin significantly reduced bacterial burden, it did not cause complete eradication even at a clinical dose. This observation may be caused by two of the behaviours we have reported in previous research^22^, namely intracellular reservoirs and bacterial biofilms on the urothelial surface, but it could also be compounded by additional factors such as phenotypic tolerance and persistence.

In the post-antibiotic era, the development of alternative therapies such as bacteriophages has gained popularity^54^. We evaluated a novel bacteriophage cocktail (LCPR1) both *in vitro* and in the 3D-UHU model, revealing distinct modes of action. Within the urine microenvironment, LCPR1 exhibited a reduced activity compared with that observed in nutrient-rich media. Similarly, Lui *et al.,* reported decreased phage efficacy in urine relative to rich media^55^. These findings underscore the importance of assessing phage performance under infection-site conditions to more accurately predict clinical efficacy. In the 3D-UHU model, LCPR1 exhibited an attenuated host cytotoxicity and a reduced pro-inflammatory response compared with untreated infections, while bacterial burden was not reduced. This suggests that the anti-inflammatory effect is unlikely to be associated with bacterial cell lysis. Instead, it is possible that LCPR1 interacted with lipopolysaccharides (LPS), thereby limiting its interaction with host receptors as shown in other studies.^56^ Transcriptomic analysis of phage-epithelial infections has revealed upregulation of anti-inflammatory mediators such as IL1R2, alongside repression of key pro-inflammatory signalling pathways including NF-κB, TNF, TLR, and JAK-STAT^57^, supporting our observations.

Compared with the use of phage or antibiotics alone, combination therapy has been reported to provide superior efficacy in suppressing bacterial populations and curbing the evolution and spread of antimicrobial resistance^58^. In the static 3D-UHU model, the dual treatment (i.e., LCPR1 + nitrofurantoin) achieved a total eradication of the planktonic bacteria, outperforming the individual treatments; however, against attached or intracellular communities, the dual treatment was not superior to nitrofurantoin alone. From an evolutionary perspective, the simultaneous action of distinct selective pressures (e.g., phage and antibiotics) may limit the capacity of bacteria to evolve resistance. Moreover, the host-phage interactions might play a crucial role in the infection dynamics.^59^

Apart from intracellular bacteria and biofilms, bacterial filaments are commonly found in UTI ^60^. Bacterial cell elongation or filamentation is thought to be a by-product of stress (including pH, UV light or antibiotics)^61^. Studies in murine UTI models have revealed that UPEC filamentation is an adaptative mechanism that promotes bacterial survival, allowing UPEC to evade or delay phagocytosis by neutrophils and macrophages^62,63^. Additionally, Aguilar-Luviano *et al.,* have shown that filaments confer protection during toxic stress by decreasing the surface area to volume ratio, which slows the diffusion and accumulation of harmful compounds within the bacterial cell^64^. In the static 3D-UHU model, we found that bacterial filamentation occurred in both UPEC and *E. coli* isolates from healthy donors in static conditions^22^. Isofidis *et al.,* characterized the filamentous IBC dispersal at a single-cell level using a microfluidic bladder cell model, and showed that the urinary environment seemed to trigger UPEC filamentation, perhaps due to acidic pH or some urine component^65^. However, the role of mechanical stimuli, such as shear stress, remains unexplored. Recently, Nadal *et al.,* show that buckling of *E. coli* filaments alters Min protein pattering and division site placement, a clear demonstration that mechanical forces and filament geometry can feed back to the cell division machinery^66^.

The urinary system has an essential and specific fluid dynamic microenvironment. Despite this, most organoid/microtissue infection work has been performed in static conditions, most likely because of the complexity and/or cost of commercially available flow-integrated systems. Introducing flow into model systems is essential to more faithfully reproduce the biophysical and biochemical cues that shape host-pathogen interactions *in vivo*. In static conditions, the absence of wall shear stress and nutrient gradients fails to capture key aspects of the infection microenvironment. To explore pathogenic strategies in the gut mucosa, Mikhaleva *et al.,* established a mucus-secreting human colonoid tissue under flow that replicated conditions experienced in the gastrointestinal tract to investigate *E. faecalis* pathogenicity^67^. In UTI, a bladder-chip model that allows cyclic stretching and flow during the filling/voiding phases was used by Sharma *et al.,* to investigate UPEC invasion and immune infiltration in real time^21^. These microfluidic platforms offer the advantage of single-cell resolution and the ability to generate finely controlled laminar flow conditions; however, they often lack the 3D complexity of the urothelium, and such systems are beyond the economic means and technical expertise of many laboratories. For the current study, we designed a relatively inexpensive mesofluidic system called P-FLO to incorporate flow into the apical side of commercially available 12-well plate Transwells, which is compatible with the 3D-UHU model and many other organoid systems. Notably, we have made these designs freely available to the community for widespread adoption across laboratories (availableava bl e on request). Although filamentation occurred in static conditions, 6 h of flow increased the frequency of UTI89 elongation in groups that were exposed to both filling/voiding cycles (with alternating low and high wall shear stress levels) or to filling only. Even at the lowest wall shear stress tested, bacterial cell elongation frequency increased in comparison with static conditions. Bacterial attachment under physiological flow was not impaired, as similar levels of bacterial coverage and bacterial burden were observed in comparison with static conditions. Apart from influencing bacterial morphology, shear stress did not noticeably alter the UPEC invasion mechanism as previously observed in mice^68^, shed patient cells^69^, human biopsies^70^ and in the 3D-UHU static infection model (here and in Flores *et al.,*^22^).

Despite the increased overall drug exposure under flow conditions, nitrofurantoin, LCPR1, and their combination did not outperform the static two-dose regimen, as comparable reductions in urothelium-associated bacteria were observed. Enhanced bacterial attachment and the presence of intracellular bacterial communities under flow may contribute to this outcome. Under static conditions, CFU data and bacterial coverage analyses were consistent; however, under flow, these measurements diverged, suggesting differences in bacterial survival and attachment dynamics. This discrepancy likely reflects physiological adaptations to shear stress, including altered metabolic states during filamentation, which increase bacterial surface area and promote adhesion. Notably, bacterial attachment under flow does not necessarily indicate viability, as adherent cells were non-viable after antibiotic exposure and dual treatment. In contrast, against planktonic bacteria, the antimicrobial effect of nitrofurantoin and the combinatorial treatment was reduced under flow compared with static conditions, likely due to enhanced nutrient exchange and cell filamentation. Together, these findings highlight the diverse influence of flow on pharmacokinetics and bacterial physiology, shaping cell elongation, attachment, and metabolic adaptation driven by altered nutrient gradients and mechanical forces.

## Conclusion

We examined the antimicrobial activity of the widely used antibiotic nitrofurantoin against UPEC across progressively complex environments. These included human urine, a static human urothelial microtissue model, and a novel mesofluidic urothelial model built from simple, low-cost 3D-printed components to create a more physiologically relevant flow-based environment. In addition, we employed both the static and mesofluidic models to evaluate the efficacy of bacteriophages alone and in combination with antibiotics. Collectively, these experiments revealed the profound influence of microenvironmental context on UPEC infection dynamics and treatment response. Although human cell-based 3D organoid, microtissue and organ-on-chip models lack crucial *in vivo* animal model components such as a blood supply, a complete repertoire of other cell types and a full immune system, they are nevertheless emerging as crucial complementary approaches to animal models, providing a human-specific micro-environment to understand host-pathogen interaction and treatment response. Such systems will be made increasingly complex in the future, allowing a more valuable window into basic infection biology as well as novel antimicrobial testing.

## Acknowledgments

We gratefully acknowledge the Engineering and Physical Sciences Research Council (EP/V026623/1) for funding. We thank the Sir Wiliam Dunn School Electron Microscopy Facility for their support in sample preparation and imaging, and Scott Hultgren, Molly Ingersoll and Meriem El Keroui for providing strains. R.O.A.J’s PhD study was funded by Beasiswa Pendidikan Indonesia (BPI), the Indonesia Endowment Fund for Education (LPDP), the Center for Higher Education Funding and Assessment (PPAPT), and the Ministry of Higher Education, Science, and Technology of Republic Indonesia. Phage work (MRJC/MEKH) was supported by the Institute for Precision Health and Leicester Drug Discovery and Diagnostics (LD3) via the institutional MRC Impact Accelerator Account (MR/X502777/1). MEKH was an Academic Clinical Lecturer funded by the NIHR. The views expressed in this publication are those of the author(s) and not necessarily those of the NIHR, NHS or the UK Department of Health and Social Care.

