## Supplementary data for "Effect of human urinary microenvironment and fluid flow on antibiotic and phage therapy efficacy against uropathogenic *Escherichia coli*"

a) Institute of Biomedical Engineering, Department of Engineering Science, University of Oxford, Oxford, UK
b) Nuffield Department of Orthopaedics, Rheumatology and Musculoskeletal Sciences (NDORMS), University of Oxford, Oxford, UK
c) Department of Renal Medicine, Division of Medicine, University College London, London, UK
d) Becky Meyer Centre for Phage Research, Division of Microbiology and Infection, University of Leicester, Leicester, UK
e) School of Biosciences, University of Sheffield, Sheffield, UK
f) Co-senior authors
*To whom correspondence should be addressed

### Supplementary Figure legends

**Figure Supplementary 1.** Dose response graphs of *E. coli* strains in caMHB (blue) and 70% pHU (yellow) against fosfomycin, trimethoprim, cefalexin and amoxicillin. The dashed pink line defines the EUCAST breakpoint. The error bar represents the standard deviation between the three biological replicates, and the empty dots represent the mean.

**Figure Supplementary 2. (A)** Schematic diagram showing the urothelium and a staining protocol to differentiate between intracellular and extracellular bacterial populations. **(B)** 3D confocal rendering of urothelium infected with *E. coli* UTI89 20 hpi, showing a cross section along the x- and y-axis to reveal intracellular bacterial communities (IBCs) within host cells, indicated with the yellow dotted line. **(C)** 3D confocal rendering and **(D)** cropped view extracted a few micrometres below the apical surface along the *z*-axis, revealing intracellular bacteria within host cells. *E. coli* UTI89 with just nucleic acid staining (cyan) indicate intracellular invasion, while extracellular bacteria are marked by combined nucleic acid staining (cyan) and anti-*E. coli* antibody (magenta). The F-actin of the eukaryotic cell is shown in purple. The letter *n* indicates nuclei **(E)** Bacterial load by CFU enumeration in the apical side (planktonic) after 12h infection, post-4h 1st treatment (Tx1) and post-4h 2nd treatment (Tx2). A Mann-Whitney test showed statistically significant difference between the untreated control and the group treated with nitrofurantoin (7xMIC) at Tx1 ($p=0.0007$, ***) and at Tx2 ($p=0.0016$,**). **(F)** Bacterial load by CFU enumeration of the attached/intracellular bacteria after the two consecutive treatments. A Mann-Whitney test was performed to compare the two groups, and no statistical difference was observed ($p=0.1419$, ns). **(G)** Dose response against nitrofurantoin in caMHB against bacterial populations recovered from the urothelium post-infection (untreated or exposed to 7xMIC). The error bar represents the standard deviation between the three biological replicates, and the empty dots represent the mean of the replicates.

**Figure Supplementary 3. (A)** Diagram showing the 3D-UHU and cropped view below the apical surface. A staining protocol was used to differentiate between intracellular and extracellular bacterial communities. *E. coli* UTI89 RFP shown in orange with nucleic acid staining (cyan) indicate intracellular invasion, while extracellular bacteria are marked by combined RFP signal (orange), nucleic acid staining (cyan), and anti-*E. coli* antibody (magenta). The F-actin of the eukaryotic cell is shown in purple. **(B)** 3D confocal rendering (left) and cropped view (right) extracted a few micrometres below the apical surface along the *z*-axis, revealing intracellular bacteria in the untreated control (**C**) and nitrofurantoin treatment (16xMIC).

**Figure Supplementary 4. (A)** Growth kinetics of *E. coli* UTI89 monitored by $OD_{600}$ over 20 h exposed to LCPR1 (MOI 1, 10 and 100) in MHB and **(B)** to 70% pHU:MHB. **(D)** CFU enumeration of *E. coli* UTI89 after 24h exposure to LCPR1 in MHB and **(E)** 70% pHU:MHB. **(C)** PFU enumeration of LCPR1 (UP17, Leic001 and JK03) after co-infection with *E. coli* UTI89 for 24h at MOI1, 10 and 100 in MHB and **(F)** 70% pHU:MHB. **(G)** PFU enumeration of LCPR1 (UP17, Leic001 and JK03) 20 hpi in the apical compartment of the 3D-UHU microtissue model infected with *E. coli* UTI89 after Tx1 **(H)** and Tx2 **(I)** collected from the 3D-UHU microtissue lysed 20phi. Uninfected 3D-UHU models exposed to LCPR1 were used as controls. All data were log-transformed to meet assumptions of normality, and differences between groups were assessed by ANOVA.

**Figure Supplementary 5. (A)** Schematic representation of the P-FLO experiment. Uninfected, fully differentiated 3D-UHU models (d≥16) were connected to the P-FLO system. pHU was flowed into the apical compartment of the

Trasnwell using a flow profile comprising 6h at 0.002 mL min$^{-1}$, interrupted every 2 h by a voiding phase of 1 min with a flow rate of 0.7 mL min$^{-1}$. **(B)** Epithelial barrier integrity measured by TEER in uninfected, fully differentiated bladder microtissue models pre and post-insertion into the P-FLO system. At least three biological experiments were performed; each dot represents a biological replicate. **(C)** Representative 3D rendering and maximum projection of confocal microscopy images of uninfected 3D-UHU in static conditions and **(D)** 3D-UHU microtissues connected to the P-FLO system: urothelial differentiation markers CK20 (orange) and UPIII (green); nucleic acid (cyan); F-actin (purple).

**Figure Supplementary 6. (A)** Experimental set-up where fully differentiated 3D-UHU (d≥16) were infected with *E. coli* UTI89-GFP at 5.0 x $10^6$ bacterial cells mL$^{-1}$, immediately after, the P-FLO system was used to apply a flow profile comprising 6 h at 0.002 mL min$^{-1}$, interrupted every 2 h by a voiding phase of 1 min with a flow rate of 0.7mL min$^{-1}$ (voiding/filling) or alternatively comprising of a 6 h exposure to 0.002 mL min$^{-1}$ flow (filling only). **(B)** Numerical simulation of the wall shear stress field in the filling phase (0.002 mL min$^{-1}$) and the voiding phase (0.7 mL min$^{-1}$), expressed in N m$^{-2}$. In the square boxes, areas of the urothelium experiencing different wall shear stress levels are indicated as high shear stress (HSS), intermediate shear stress (ISS), and low shear stress (LSS). **(C)** Representative 3D rendering of image of *E. coli* UTI89-GFP 6 hpi in the 3D-UHU using the filling protocol. UTI89-GFP is represented in orange, the nucleic acid staining in cyan, and the cytoskeleton staining in purple. **(D)** Percentage bacterial area coverage on top of the urothelium obtained from confocal imaging. Data are plotted as means (line) and each empty dot represents a technical replicate (different images) from the three biological experiments. An ordinary Anova test was performed to compare the different groups. **(E)** Cumulative fraction plot of the bacterial length obtained under filling-only and voiding/cycle regimens and grouped into HSS, ISS and LSS. **(F)** Representative maximum projection confocal obtained in the HSS, ISS and LSS regions.

**Figure Supplementary 7. (A)** Schematic diagram showing the urothelium and a cropped version in the z-axis to reveal IBCs. (**B)** 3D confocal rendering of urothelium infected with *E. coli* UTI89 20 hpi under flow and **(C)** a 3D cropped view extracted a few micrometres below the apical surface along the *z*-axis to reveal IBCs. **(D)** Top view of a representative 3D rendering confocal microscopy image of *E. coli* UTI89 untreated at 20 hpi under static conditions and **(E)** under flow. *E. coli* UTI89-GFP is shown in orange, nucleic acid staining in cyan and actin in purple. **(F)** Comparison of the log reduction caused by antimicrobial treatments against the planktonic bacteria and **(I)** the urothelium-associated bacterial population, under static and flow conditions. **(G)** PCU enumeration of LCPR1 (UP17Leic001 and JK03) in the apical compartment of the 3D-UHU model infected *E. coli* UTI89 20 hpi under flow and **H)** collected from the lysed microtissue comportment. The phage recovery data were log-transformed to meet assumptions of normality, and differences between groups were assessed by Anova.

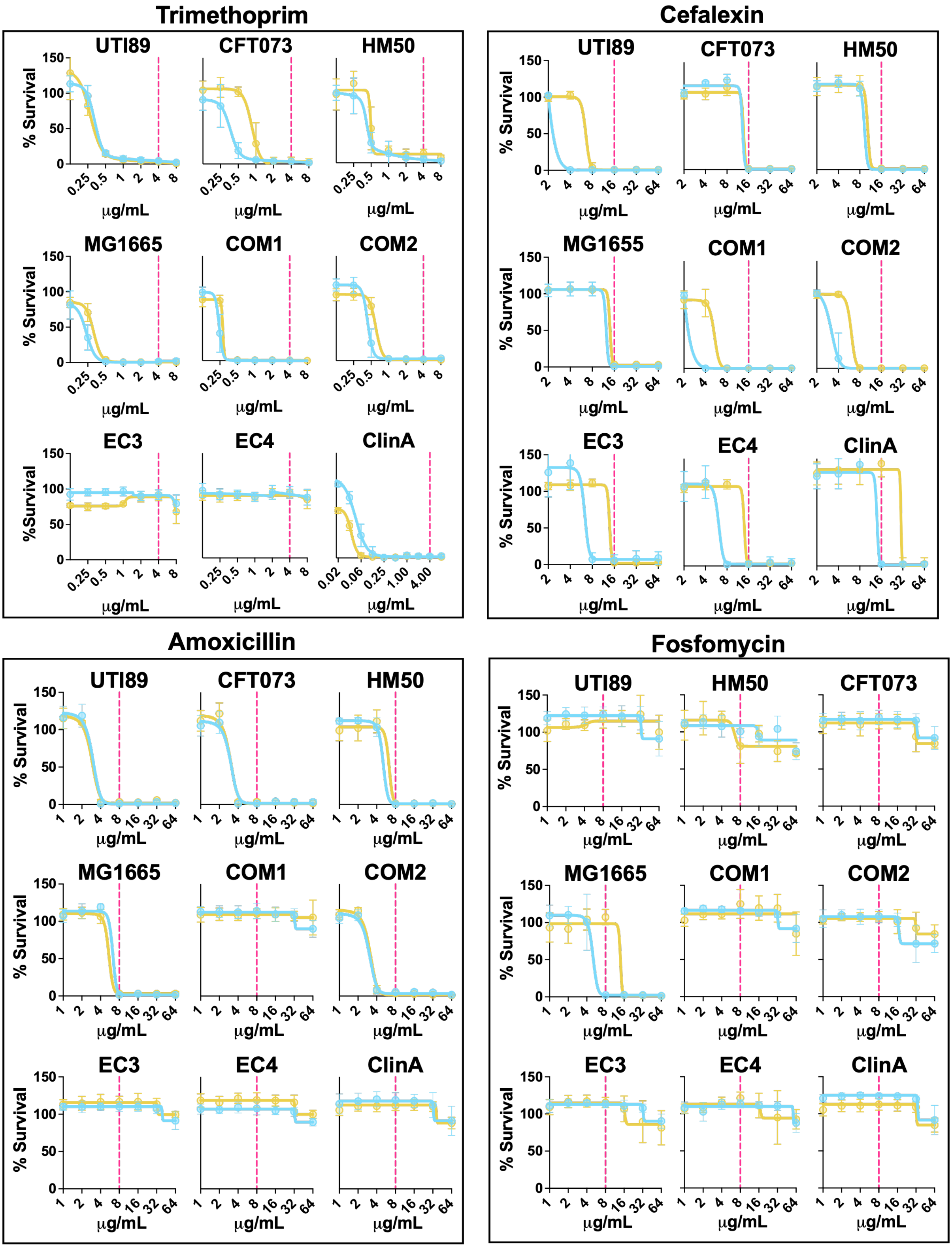


**Figure Supplementary 1.**

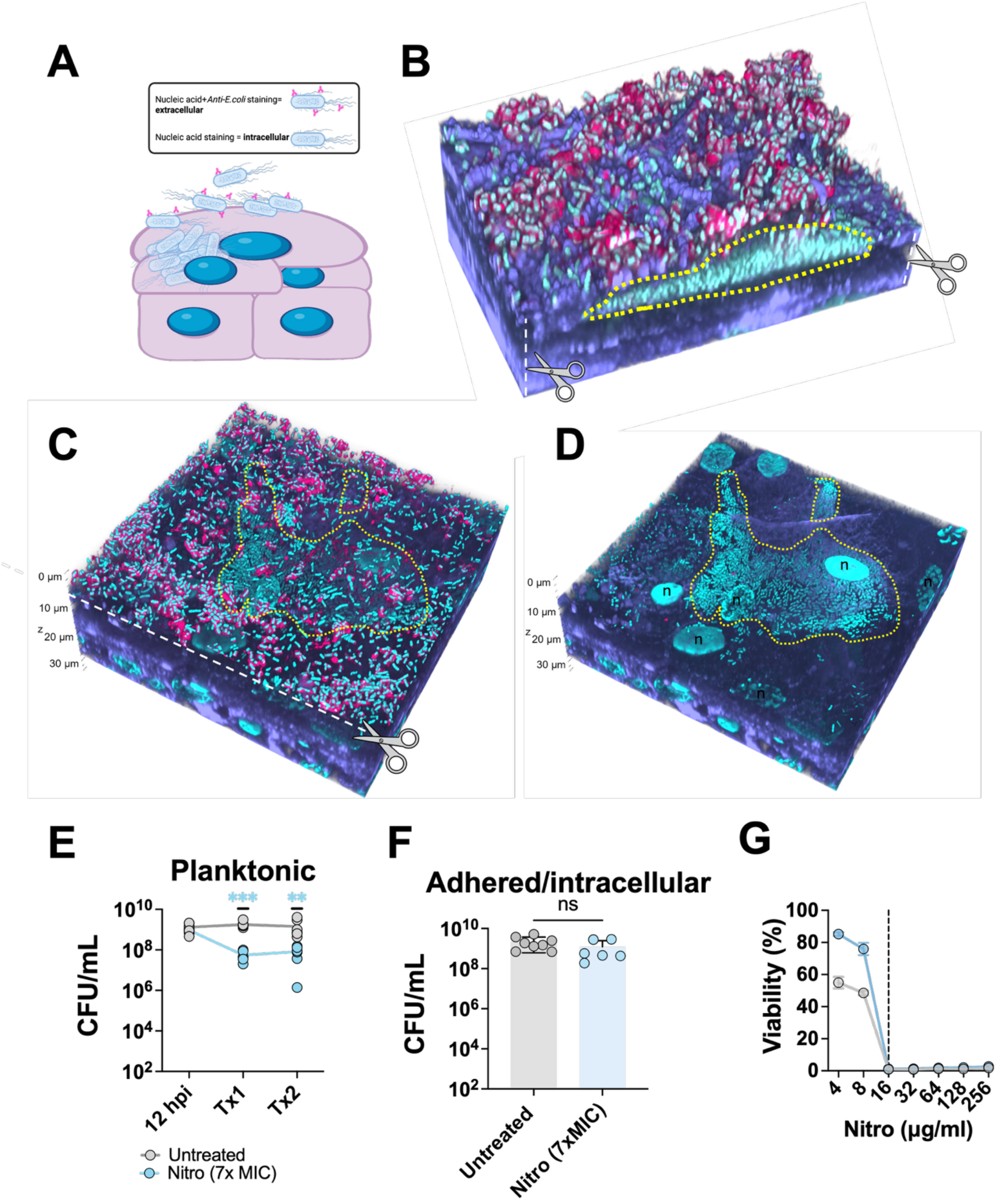


**Figure Supplementary 2.**

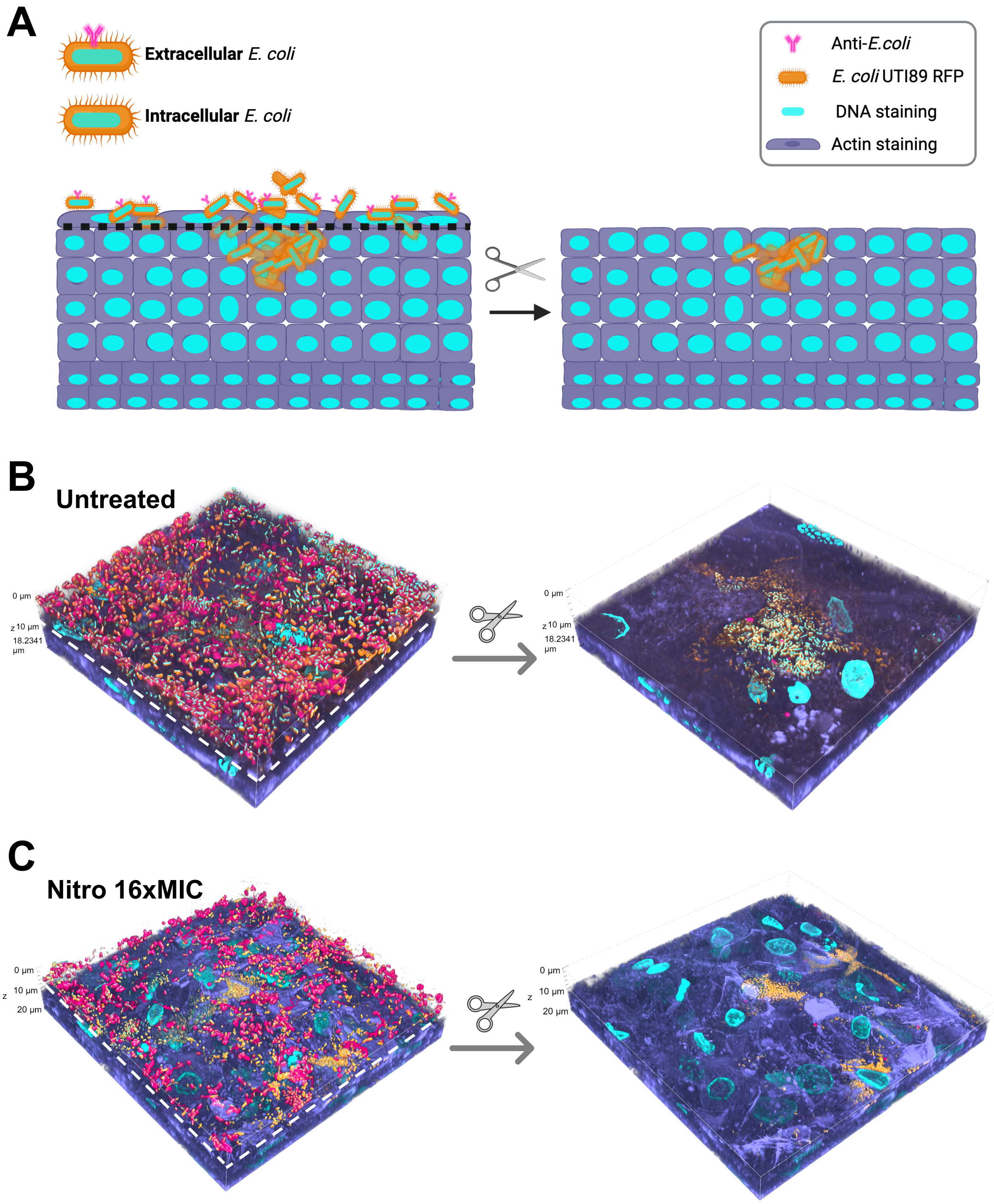


**Figure Supplementary 3.**

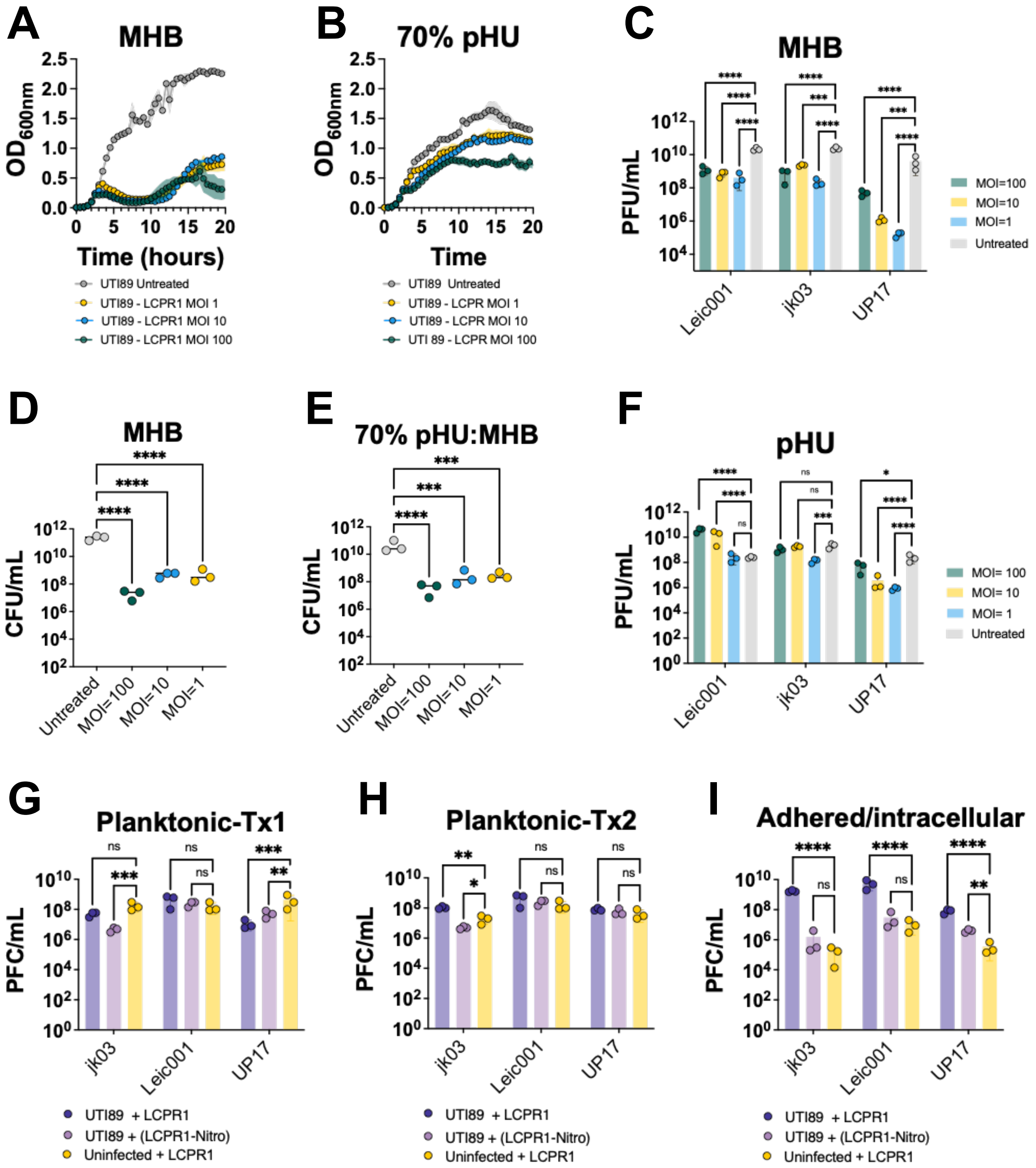


**Figure Supplementary 4**

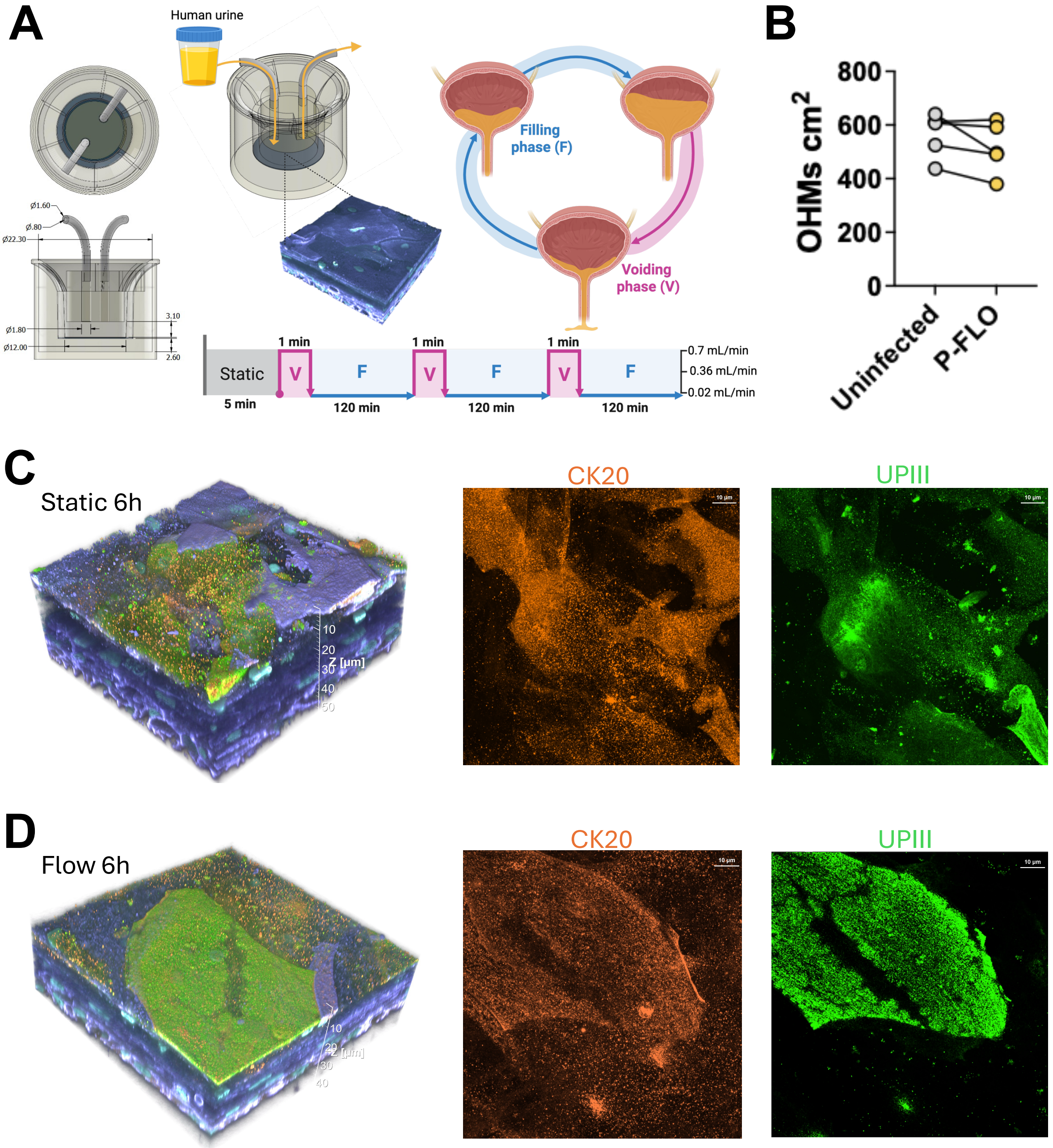


**Figure Supplementary 5.**

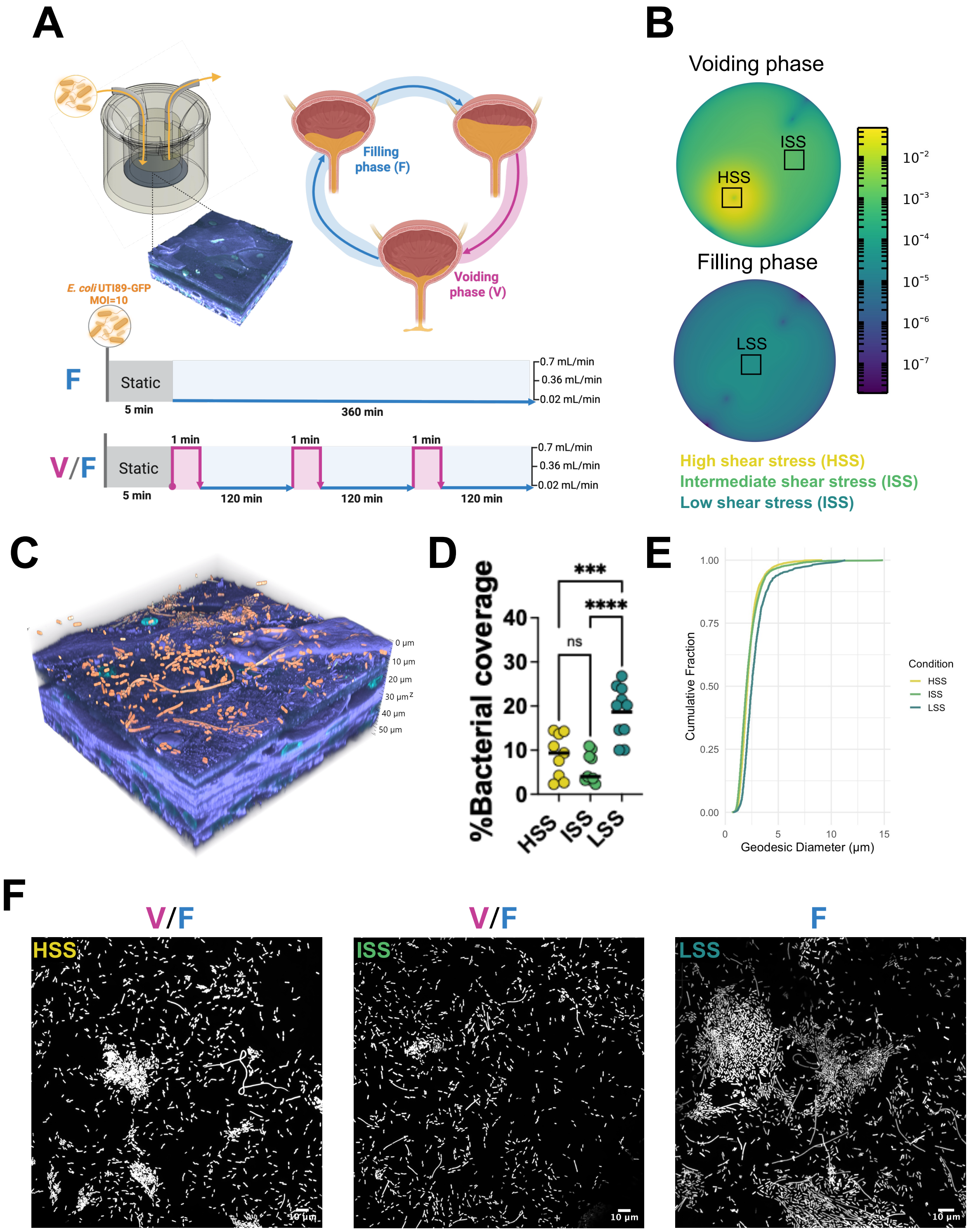


**Figure Supplementary 6.**

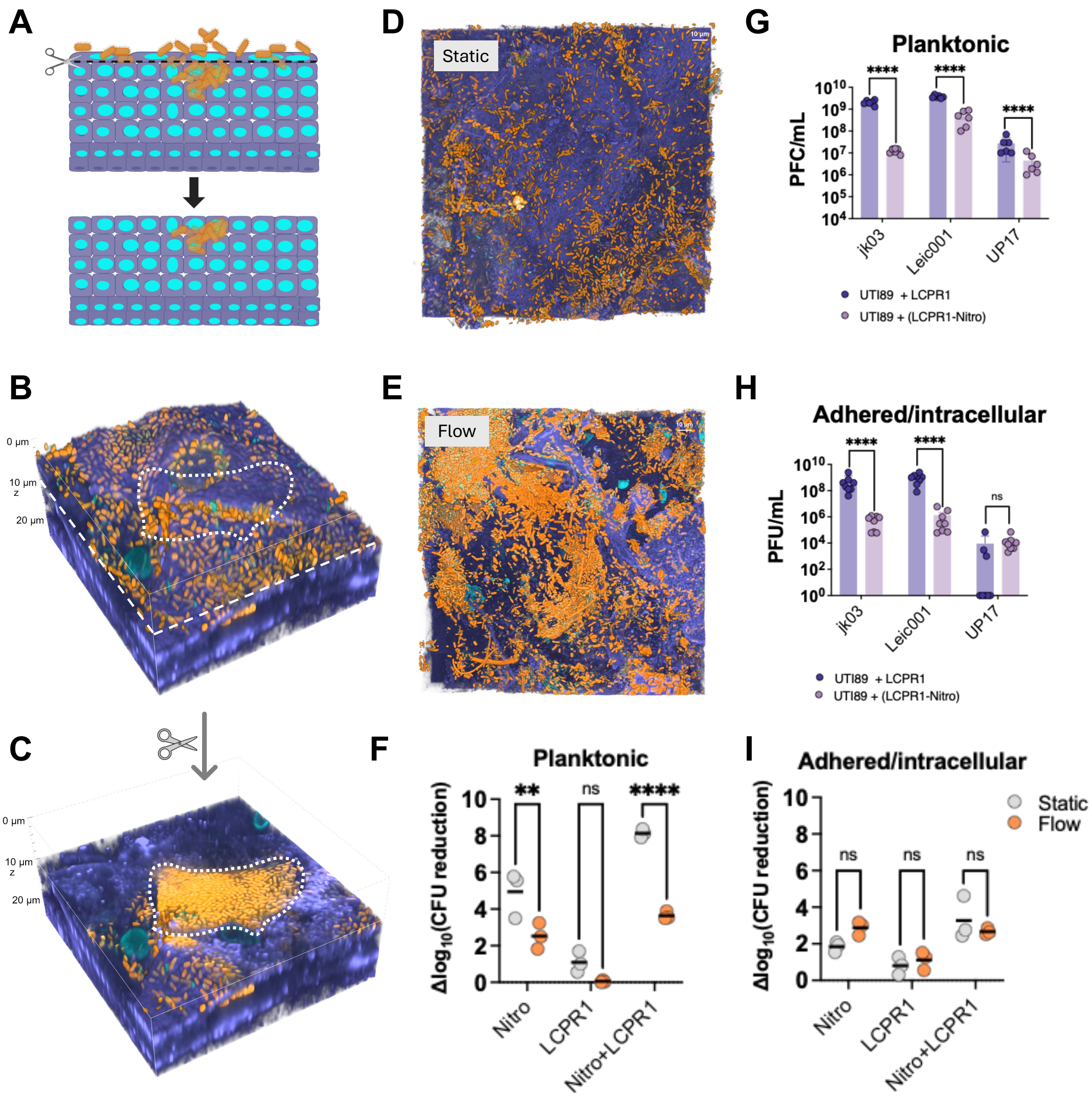


**Figure Supplementary 7.**

**List of phages used in this study (LCPR1)**

| Phage Name | TEM | Genus | Genome Size | Reference |
|---|---|---|---|---|
| **vB_KpnM_311F** | 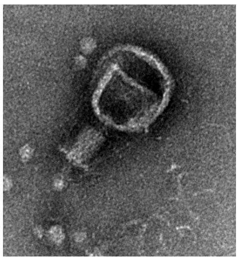 | ***Jiaodavirus*** | **167 kb** | 71 |
| **vB_KpnM_05F** | 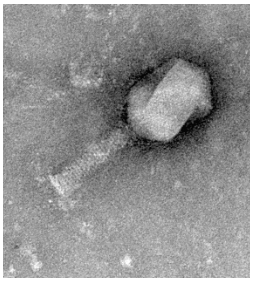 | ***Jiaodavirus*** | **175 kb** | 71 |
| **vB_KpM_Centimanus (Phage 71)** | 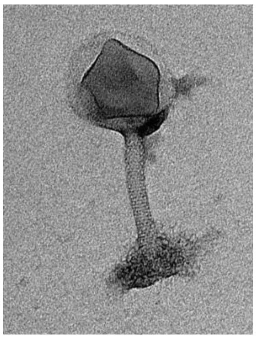 | ***Maaswegvirus*** | **299 kb** | **-** |
| **vB_KppS_Storm (Phage 34)** | 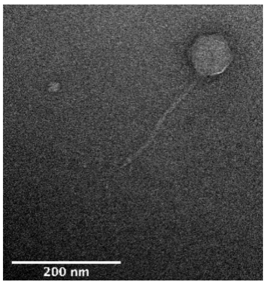  | ***Sugarlandvirus*** | **110 kb** | 72 |
| **vB_KppS_Anoxic (Phage 52)** | 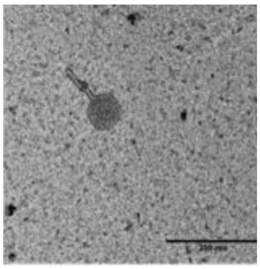 | ***Sugarlandvirus*** | **109 kb** | 72 |

| | | | | |
|---|---|---|---|---|
| **vB_KpM_SoFaint (Phage 70)** | 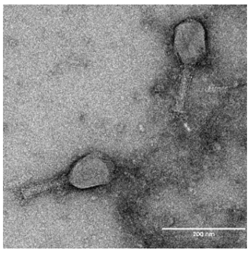 | ***Slopekvirus*** | **176 kb** | [72] |
| ***E.coli*** **Phage** | | | | |
| **vB_EcoM_UP17** | 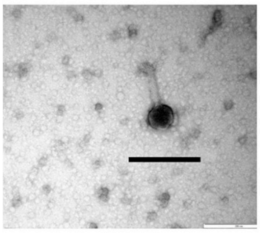 | ***Myovirus*** | **152 kb** | [71] |
| **JK03** | 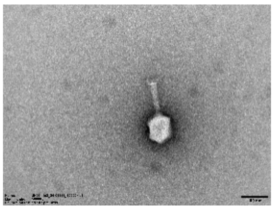 | ***Myovirus*** | **170 kb** | **-** |
| **Leic_001** | 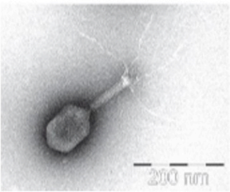 | ***Myovirus*** | **169 kb** | **-** |
